# Aminoacyl-tRNA synthetase organization is crucial for mammalian brain development

**DOI:** 10.1101/2025.09.08.674979

**Authors:** Samuel Protais Nyandwi, Rasangi Tennakoon, Alana Hoffmann, Yong Jia Bu, Qingyu Shi, Sofija Volkanoska, Danyon Harkins, Flavia de Lima Alves, Colette Maya Macarios, Sascha Woolcott, Jingchun Zhou, Han Gao, Jiachen Cai, Edward Devereux, Aaron R. Wheeler, Mark Nitz, Veronique E. Miron, Scott A. Yuzwa, Haissi Cui

## Abstract

Central nervous system development relies on spatially regulated protein synthesis, which includes the aminoacylation of tRNAs. Developmental brain disorders, such as leukodystrophies, can be caused by the exclusion of Arginyl-tRNA synthetase (RARS1) from the multi-tRNA synthetase complex, which does not affect its enzymatic function.

To understand the purpose of aminoacyl-tRNA synthetase complex formation, we engineered a mouse line where RARS1 is untethered from the complex in forebrains. We observed profound behavioral phenotypes and anatomical changes originating from a severe reduction in oligodendrocytes and subsequent myelination defects. On the molecular level, disruption of the multi-tRNA synthetase complex caused RARS1 mis-trafficking, which resulted in lost colocalization with ribosomes, leading to the concomitant activation of integrated stress response. Proteomics analysis of newly synthesized proteins suggested that mRNA translation in cellular projections was specifically reduced, affecting myelin structural proteins.

In summary, exclusion of RARS1 from the multisynthetase complex led to its altered subcellular localization and decreased protein synthesis in oligodendrocyte projections, directly linking molecular effects to white matter pathologies. Altogether, our results reveal the necessity of aminoacylation organization for mammalian neurodevelopment.

## Introduction

Leukodystrophies are neurological disorders caused by defective myelination^1–3^, a developmentally regulated process whereby oligodendrocytes wrap lipid membranes (myelin) along axons for faster signal conduction. Myelination is inherently linked to oligodendrocyte differentiation^2,4–6^ and a lack of myelination subsequently causes neuronal death^4,6^. While the genetic origins of myelination defects are highly heterogeneous, mutations in genes associated with protein synthesis are specifically associated with insufficient myelination, termed hypomyelination^2,3^.

Aminoacyl-tRNA synthetases (ARSs) are crucial for protein synthesis due to their role in charging tRNAs with their cognate amino acids, thereby enabling accurate mRNA translation^7,8^. Although aminoacyl-tRNA synthetases are ubiquitously expressed, disorders caused by mutations in their genes are predominantly linked to neurodevelopmental disorders^2,9^. In mammals, 9 aminoacyl-tRNA synthetases together with three designated adapter proteins (AIMP1-3), form the multi-tRNA synthetase complex (MSC)^10–13^. Aminoacyl-tRNA synthetases are integrated into the MSC either through designated domains or by interactions mediated by their catalytic domains^12,14–17,8^. Despite an evolutionary drive towards complex formation^12^, the role of the MSC is still unclear: it has been linked to improved protein synthesis by channeling charged tRNAs to ribosomes^18–20^, but exclusion of individual ARSs, especially Arginyl-tRNA synthetase (RARS1), does not disrupt bulk protein synthesis and Arg decoding^21^. The MSC is also associated with noncanonical functions^8,22,23^, such as regulating the release of individual aminoacyl-tRNA synthetases^24,25^.

Mutations in RARS1^26–28^ are causal for hypomyelinating leukodystrophies, a heterogeneous group of white matter disorders characterized by a lack of myelin deposition. Interestingly, the majority of mutations found in RARS1^28^ cause its exclusion from the MSC: RARS1 is anchored to the MSC by its N-terminal coiled-coils, which are referred to as leucine zippers (LZ)^15,21,29^. Of note, the RARS1 N-terminal coiled-coil domain does not formally fulfill the definition for leucine zippers^30^, but we will refer to it as such in line with previous publications^23,31^. Leukodystrophy-associated mutations result in the preferential translation from a second start codon, thereby altering the ratio between complex-bound and N-terminally truncated, free RARS1^28,32^. This connection strongly suggests that MSC formation is specifically important for brain development.

To understand the importance of the ARS complex formation, we excluded RARS1 from the MSC in the forebrain of mice. We observed phenotypical changes similar to those in patients with leukodystrophy, namely microcephaly and reduced myelination. RARS1’s subcellular localization was severely disrupted, highlighting a connection between MSC formation and RARS1 trafficking. Reduced localized mRNA translation could be directly linked to pathological changes. Altogether, these findings suggest MSC formation is crucial for mammalian brain development and specialized protein synthesis.

## Results

### Generation of a mouse model to investigate RARS1-driven pathology

To study the influence of RARS1 on neural development, we generated a mouse line in which exon 2 of RARS1, which encodes the majority of the RARS1 LZ, was flanked by loxP sites (**Figure 1A**). Upon introduction of an appropriate Cre driver, recombination causes the deletion of exon 2, leading to an in-frame exclusion of most of the leucine zipper coding region (45 of 72 amino acids). We chose this design instead of the point mutations found in patients for two reasons: first, mutations in the start codon or unproductive translation from the first start codon significantly lowered RARS1 protein levels^21^, making it difficult to distinguish between effects caused by the reduction of total RARS1 aminoacylation capacity in cells and altered complex integration of RARS1. Second, the deletion of an exon is more easily controlled by Cre-drivers and allows for tissue-specific RARS1 LZ deletion. Homozygous systemic truncation of RARS1 by crossing *Sox2*-Cre mice^33^ with *Rars1^loxP/loxP^* mice led to premature death at P0 due to a lung defect (pathology report IDEXX) (**Extended Data Fig. 1A-C**). As myelination in mice starts after birth^34^, we opted to limit the deletion of exon 2 to the brain. We chose *Emx1*-controlled Cre expression^35^, where the RARS1 LZ is deleted in excitatory neurons in the forebrain as well as in glial cells of the pallium early in development, before E10^35^ (**Figure 1A**). Cre-positive animals homozygous for *Rars1^loxP/loxP^* were obtained at wean age, albeit at sub-Mendelian ratios, at equal distribution of sexes (**Figure 1B**). Tissue specific deletion of the RARS1 LZ in the telencephalon thus circumvented lethality while allowing us to observe effects throughout neurodevelopment.

**Figure 1:**
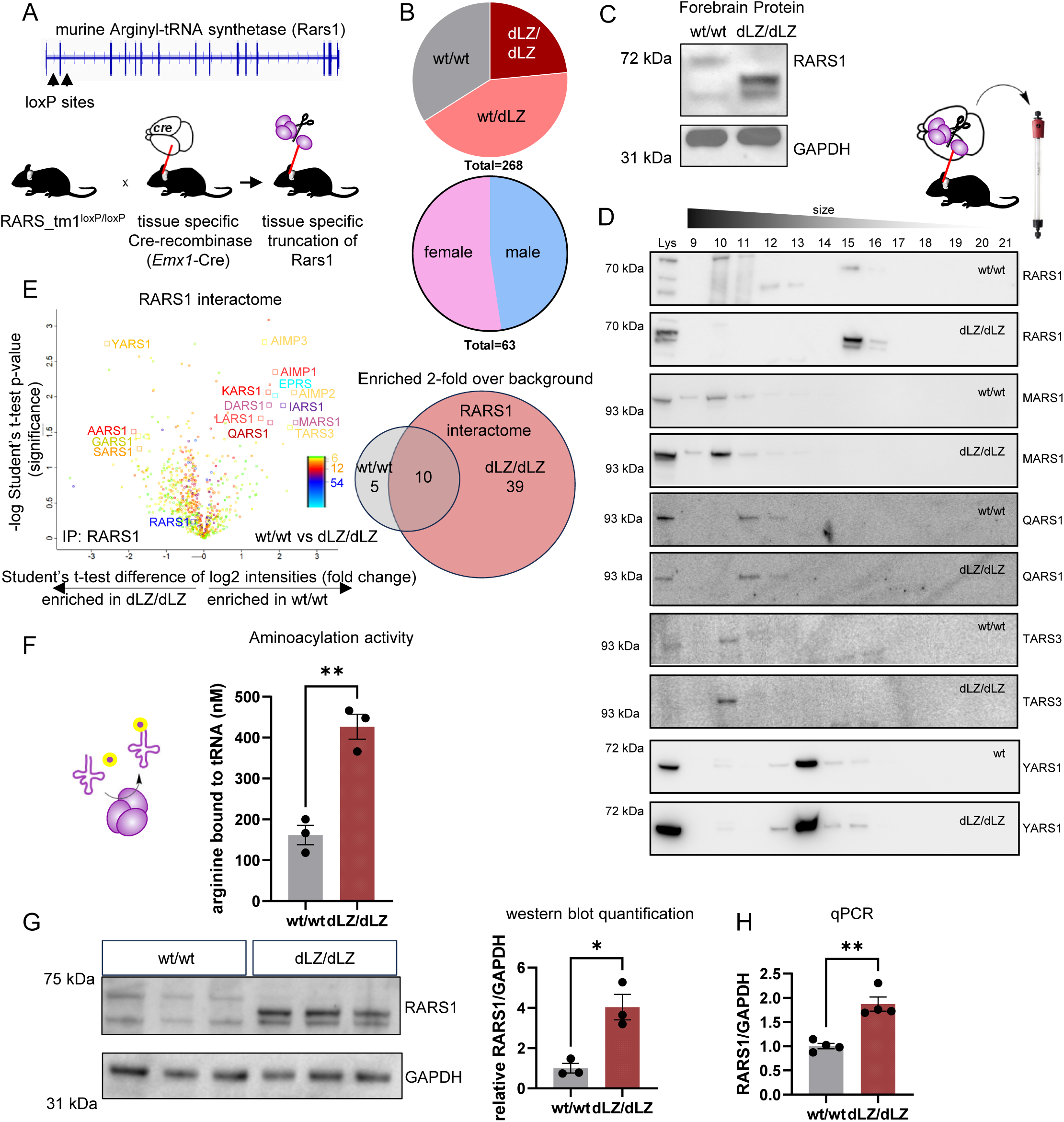
Deletion of the RARS1 leucine zipper causes its exclusion from the multisynthetase complex but does not reduce enzyme levels and activity. A) Top panel: Design of a mouse line, where RARS1 can be N-terminally truncated by deletion of exon 2 (flanked with loxP sites). Bottom panel: Breeding scheme to yield animals lacking the RARS1 leucine zipper (dLZ) in the forebrain and hippocampus. B) Genotype distribution among *Emx1-*Cre+ animals. Sex distribution among dLZ/dLZ animals. C) Immunoblot of forebrain lysate, probed for RARS1. GAPDH was used as a loading control. D) Size exclusion chromatography followed by immunoblot to assess the integrity of the multisynthetase complex. Earlier fractions correspond to higher molecular weight. MARS1, QARS1: multisynthetase complex-bound aminoacyl-tRNA synthetases. TARS3: peripheral multisynthetase complex aminoacyl-tRNA synthetase. YARS1: free-standing, dimeric aminoacyl-tRNA synthetase. E) Volcano plot of proteins identified by mass spectrometry following co-immunoprecipitation. *Y*-axis: significance (*p* value). *X*-axis: difference log_2_(fold change). Color coding denotes the number of peptides per protein. Comparison between wt/wt and dLZ/dLZ interactomes retrieved through anti-RARS1 antibody pulldowns in wt/wt and dLZ/dLZ lysates. Protein enriched more significantly in wt RARS1 pulldowns than dLZ RARS1 appear on the *left side* of the volcano plot. n = 3, 3. Student’s t-test. F) Aminoacylation activity assessed by filter binding using in vitro transcribed tRNA and radiolabeled arginine. Activity is shown after 15 minutes incubation at room temperature. n=3, 3. Unpaired, two-tailed t test, **p=0.0024. G) Immunoblot probing RARS1 protein levels and quantification of density. GAPDH was used as a loading control. n=3, 3. *p=0.0108. H) mRNA levels of RARS1 assessed using qPCR. n=4, 4. Unpaired, two-tailed t test, **p=0.0014 B-H) wt/wt: wildtype RARS1, *Emx1-*Cre+ x Rars1^wt/wt^. dLZ/dLZ: RARS1 delta leucine zipper, *Emx1-*Cre+ x Rars1^loxP/loxP^. C, D) A representative of at least triplicates is shown. C-H) Replicates are individual animals. All experiments were performed in 2 months old male mice.

Immunoblots showed almost exclusive expression of truncated RARS1 at the expected molecular weight in the forebrain (**Figure 1C**) but, as expected, not in the spleen (**Extended Data Fig. 1D**). We will subsequently refer to *Cre+/Rars1^wt/wt^* mice as wt/wt, *Cre+/Rars1^wt/loxP^* mice as wt/dLZ and homozygous *Cre+/Rars1^loxP/loxP^*mice as dLZ/dLZ. The naturally occurring shorter form of RARS1^29^ was also observed in both wt/wt and dLZ/dLZ animals (**Figure 1C**), in addition to the Cre-induced truncation.

In order to confirm that RARS1 was lost from the MSC, we performed size exclusion chromatography to separate free, unbound aminoacyl-tRNA synthetases from their MSC-bound counterparts^21,36^. We lysed mouse forebrains under mild conditions to ensure complex integrity and identified aminoacyl-tRNA synthetases using immunoblot (**Figure 1D**, quantification in **Extended Data Fig. 1E**). ARS associated with the MSC, such as methionyl-tRNA synthetase (MARS1) and wt RARS1, eluted at fractions 9-11, while free aminoacyl-tRNA synthetases, such as Tyrosyl-tRNA synthetase (YARS1), corresponded to fractions 13-16. RARS1 in wt animals predominantly eluted at fraction 10-11, indicating its complex bound form, but free RARS1 could also be found in fraction 15. In contrast, in dLZ/dLZ mice, RARS1 was exclusively found in fraction 15 and 16. Two bands could be seen in these fractions, indicating protein lacking exon 2 encoded amino acids as well as the endogenous leucine zipper-free form. In line with previous reports^17,37^, MARS1 was still found at higher molecular weight fractions, supporting ARS complex formation in the absence of RARS1^21,38^. Free-standing ARS, such as YARS1, were also not affected by RARS1 exclusion and eluted at the same molecular weight in both wt/wt and dLZ/dLZ lysates.

Removal of RARS1 from the multisynthetase complex in a cell line model led to the simultaneous exclusion of QARS1^21^. Interestingly, in forebrain lysates, QARS1 eluted in fractions 9-15 in both wt/wt and dLZ/dLZ mice, indicating that QARS1 was retained (**Figure 1D**, alternative antibody in **Extended Data Fig. 1F**). We therefore probed TARS3 (ThrRS-L protein), which contains a similar LZ as RARS1^39^. TARS3 eluted in fraction 10 in both wt/wt and dLZ/dLZ lysates, corresponding to its integration into the MSC. This suggested that TARS3 can substitute for RARS1 to anchor QARS1 in the complex.

We further investigated RARS1 protein interaction partners by co-immunoprecipitation mass spectrometry in wt and dLZ/dLZ animals. We found MSC components between ∼3- to ∼5.5-fold less enriched in dLZ/dLZ animals, confirming the loss of RARS1 from the MSC (**Figure 1E, Supplementary Tables 1-3**). MSC components were, however, co-immunoprecipitated in both wt and dLZ/dLZ lysates, possibly due to the presence of some full-length RARS1 in dLZ/dLZ animals (**Extended Data Fig. 1G**).

### Deletion of the leucine zipper does not reduce RARS1 bulk enzymatic activity

To test whether exclusion from the MSC would affect RARS1 activity, we isolated forebrain lysates and tested their ability to aminoacylate *in vitro* transcribed tRNAs with tritium-labeled arginine^40^. Interestingly, we found that activity was increased in dLZ/dLZ forebrains compared to wt/wt (**Figure 1F, Extended Data Fig. 1H**). To understand this elevated activity, we compared protein levels of RARS1 between wt/wt and dLZ/dLZ and found that deletion of the leucine zipper led to a more than three-fold increase of RARS1 in forebrains (**Figure 1G**). Furthermore, RARS1 was upregulated at the mRNA level (**Figure 1H**), suggesting an increase of RARS1 protein due to elevated RARS1 gene expression. Together, this data confirms the exclusion of RARS1 from the MSC in mouse forebrains without reducing RARS1 bulk enzymatic activity.

### Complex behavioral abnormalities and reduced body weight in mice lacking RARS1 in the multisynthetase complex

Patients with RARS1 variants affecting the first and second codons show mild to severe symptoms, including impaired motor control, depending on the specific variant^28^. We were therefore interested in characterizing behavioral consequences of the RARS1 LZ deletion. We measured the body weight of mice at 2 months of age and found that dLZ/dLZ mice of both sexes were significantly lighter (**Figure 2A**). This weight difference was reduced by providing food on the ground and was lost at 6 months of age (**Extended Data Fig. 2A, B**), suggesting that it was predominantly caused by motor issues instead of metabolic causes.

**Figure 2:**
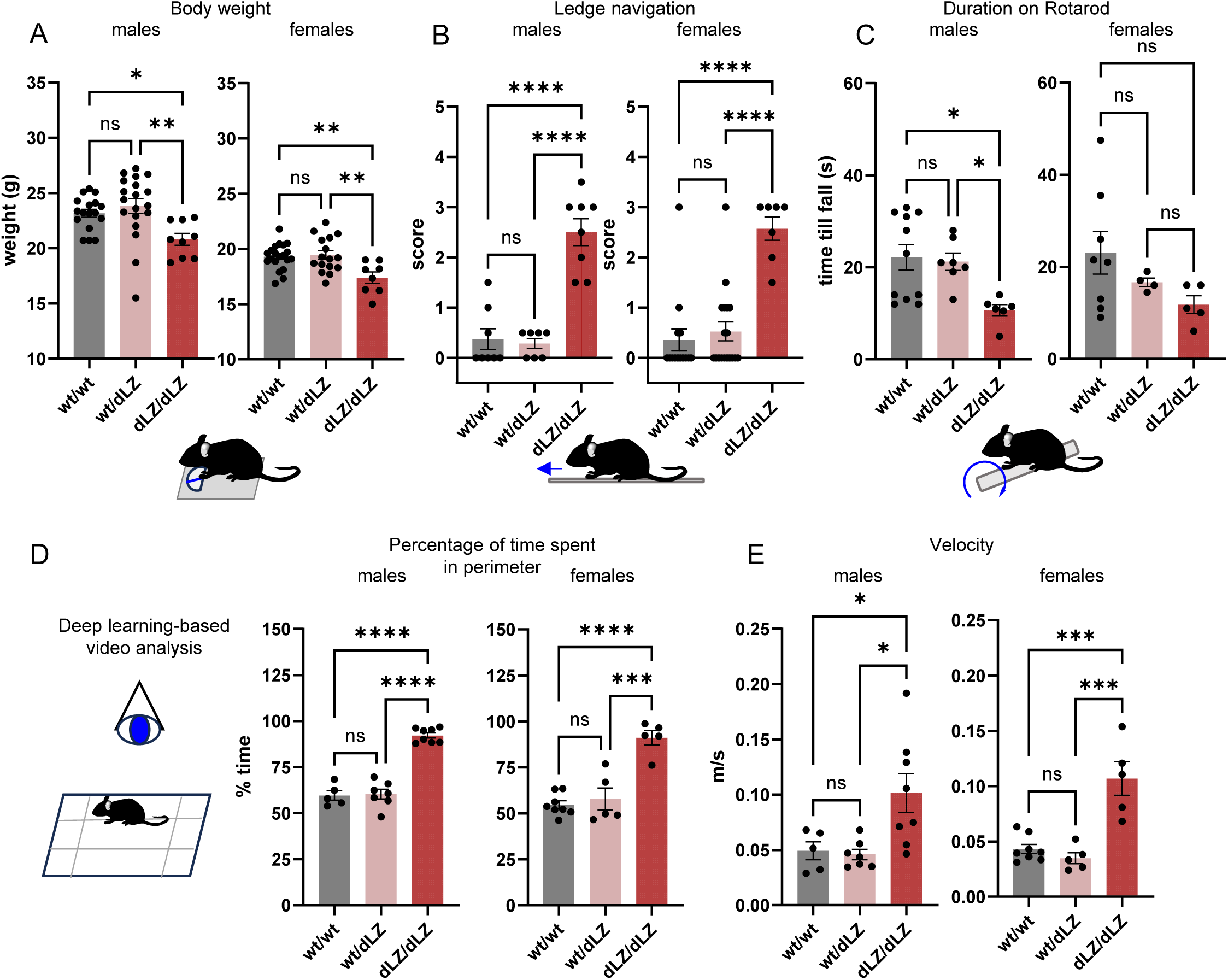
N-terminal truncation of RARS1 in the forebrain and hippocampus leads to phenotypical and behavioral changes. A) Body weight at 2 months. Males: n=17, 19, 9. Ordinary, one-way ANOVA, multiple comparisons, *p=0.0388, **p=0.0053. Females: n=20, 16, 8. Ordinary, one-way ANOVA, multiple comparisons, **p=0.0056, **p=0.0037. B) Ledge navigation: animals were placed on the ledge of a fresh cage and monitored while lowering themselves back into the cage. The higher the score, the worse the animal performed. Balance, mobility, and ability to lower themselves without assistance were scored. Males: n=8, 7, 8. Ordinary, one-way ANOVA, multiple comparisons, ****p<0.0001. Females: n=14, 19, 7. Ordinary, one-way ANOVA, multiple comparisons. ****p<0.0001. C) Rotarod: Animals were placed on a rotating rod and the speed of rotation was increased from 4 rpm to 60 rpm over 60 seconds. The time to fall is shown, corresponding to a defined rotational speed. Males: n=11, 7, 6. Ordinary, one-way ANOVA, multiple comparisons, *p=0.0109, *p=0.0350. Females: n=8, 4, 5. Ordinary, one-way ANOVA, multiple comparisons, ns p>0.05. D, E) Open field test: Mice were placed in a transparent box lined with grid paper and their behavior was filmed. D) Percentage of time spent in the perimeters of the box as opposed to the center. Males: n=5, 7, 8. Ordinary, one-way ANOVA, multiple comparisons, ****p<0.0001. Females: n=8, 5, 5. Ordinary, one-way ANOVA, multiple comparisons, ***p=0.0001, ***p=0.0001. E) Velocity as measured in m/s. Males: n=5, 7, 8. Ordinary, one-way ANOVA, multiple comparisons, *p=0.0370, *p=0.0142. Females: n=8, 5, 5. Ordinary, one-way ANOVA, multiple comparisons, ***p=0.0001, ***p=0.0001 (A-E) wt/wt: wildtype RARS1, *Emx1-*Cre+ x Rars1^wt/wt^. dLZ/dLZ: RARS1 delta leucine zipper, *Emx1-*Cre+ x RARS1^loxP/loxP^. Replicates are individual animals. All experiments were performed in 2 months old male mice.

To assess potential behavioral phenotypes and guide further studies, we performed a battery of non-invasive assays, namely ledge navigation, grip strength, hindlimb extension, and stride length assessments^41^. dLZ/dLZ male and female mice scored worse in ledge navigation (balancing on a thin cage ledge and lowering themselves back into the cage) (**Figure 2B**) but performed comparable to wt/wt littermates in all other assays (**Extended Data Fig. 2C-F**). To obtain more quantitative results, we assessed motor function using the rotarod method^42^. Male dLZ/dLZ animals performed significantly worse in rotarod (**Figure 2C**), despite having comparable grip strength to wt animals (**Extended Data Fig. 2C**), suggesting that their reduced time till fall was not due to muscle weakness. Female mice on the other hand did not show reduced rotarod performance which could be due to their overall lower body weight (**Figure 2A, C**).

Forebrain defects can alter mouse behavior in an open field^43,44^. We tested mouse movement patterns, velocity, and preference for the perimeter versus the middle of the field using a deep learning-based video analysis software^45^. Male and female dLZ/dLZ mice spent significantly more time in the perimeter of the open field and moved faster in an unknown environment, as shown by heightened velocity (**Figure 2D, E, Extended Data Fig. 2G, H**). In summary, this data suggests that motor coordination and balance issues did not arise from muscle weakness or an inability to move but rather lowered fine motor control and possibly anxiety-induced behavior.

### Reduction of forebrain volume in animals with RARS1 truncation

To investigate neurological consequences of RARS1 exclusion from the MSC, we investigated gross morphological changes in the forebrains of animals with dLZ/dLZ RARS1. Side by side alignment of wt and dLZ/dLZ brains revealed a reduction of brain size in dLZ/dLZ animals (**Figure 3A, Extended Data Fig. 3A**). Disruption in anterior-posterior brain arealization was observed in dLZ/dLZ animals; forebrain length was reduced to ∼73% of wt animals in two-month-old male mice (**Figure 3A**) and ∼83% in two-month-old female mice **(Extended Data Fig. 3A)**. The reduction remained present in six months old animals (**Extended Data Fig. 3B)**. We further assessed this striking difference by quantifying the area of depth-matched coronal sections of the forebrain (sections between Bregma +0.86 mm and Bregma +0.14 mm) (**Figure 3B, C, Extended Data Fig. 3C, D,** additional images in **Extended Data Fig. 4**). The section area was 40% smaller in dLZ/dLZ brains and a significant decrease in both cortical and corpus callosum thickness was observed.

**Figure 3:**
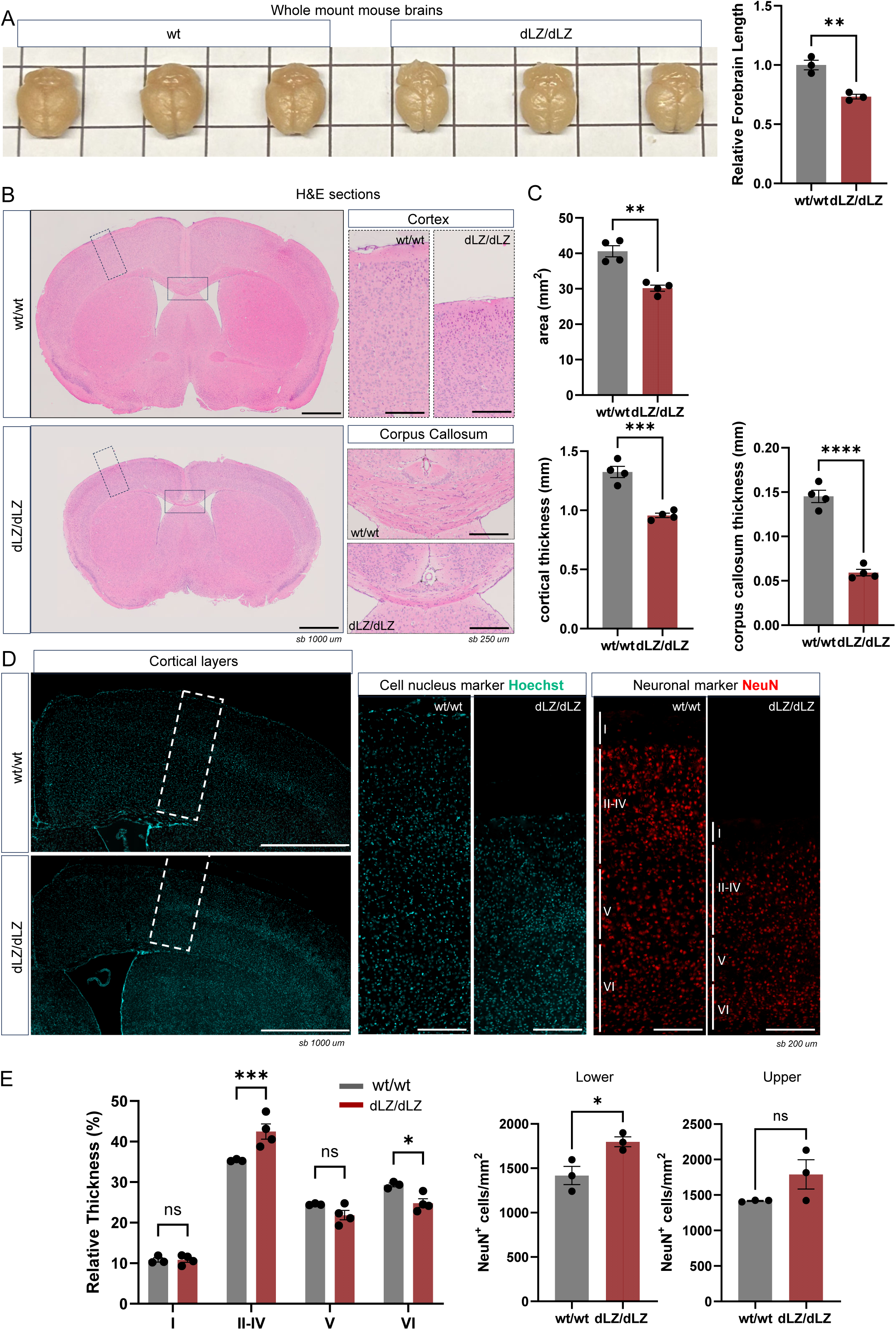
N-terminal truncation of RARS1 leads to reduced brain volume. A) Picture of whole mount brains. Grid size is 1 cm. n=3, 3. Unpaired, two-tailed t test, **p=0.0040. B) Representative FFPE sections of mouse brains stained with hematoxylin and eosin. Insets of the corpus callosum and cortex are shown. Scale Bar = 1 mm. C) Brain section area, cortical thickness along the midline, and corpus callosum thickness quantified from matched serial sections of forebrains. n=4, 4. Unpaired, two-tailed t test, **p = 0.0011, ***p < 0.0003, ****p < 0.0001. D) Neuronal marker NeuN (Rbfox3) immunostaining. Bar = 20 µm. E) Quantification of relative cortical layer thickness. Layers I-IV are superficial layers. Layer V-VI are deep layers. n = 3, 4. Two-way ANOVA, ***p = 0.0006, *p = 0.0315. F) Quantification of neuronal marker NeuN-positive nuclei. n=3, 3. Unpaired, two-tailed t test, *p = 0.0313, ns p>0.05. A-F) wt/wt: wildtype RARS1, *Emx1-*Cre+ x Rars1^wt/wt^. dLZ/dLZ: RARS1 delta leucine zipper, *Emx1-*Cre+ x Rars1^loxP/loxP^. Replicates are individual animals. All experiments were performed in 2 months old male mice. B, D) A representative image of 4 replicates is shown.

Given the decreased cortical thickness, we assessed whether cortical layering was changed in dLZ/dLZ brains to identify potential defects in neurogenesis and neuronal migration. By analyzing both the pan-neuronal marker NeuN and nuclear marker Hoechst (**Figure 3D, E**), we observed that superficial cortical layers (II-IV) were relatively thicker in dLZ/dLZ brains. We also assessed neuronal density and found that it increased in lower layers (V-VI). These findings align with reports of microcephaly in some human patients with RARS1 N-terminal truncation^28^.

### RARS1 truncation impairs myelination

To assess the underlying cause of microcephaly, we performed transcriptome analysis through RNAseq on the forebrains of 2-months old male mice. We could confirm the exclusion of exon 2 as reads mapping to junctions between exon 1-3 were vastly different between wt/wt and dLZ/dLZ tissues (**Extended Data Fig. 5A**). Strikingly, over 4000 genes were differentially expressed on the mRNA level (adjusted *p*-value < 0.05) (**Figure 4A, Supplementary Table 4**) and genotypes clearly segregated in principal component analysis (**Extended Data Fig. 5B**). Among these genes, enrichment analysis suggested that mRNAs associated with oligodendrocyte differentiation were downregulated, including Sox10 and Myrf (**Figure 4B**). As these aligned with the severe phenotypical observations on brain anatomy and reports of RARS1 variants in leukodystrophies^2,28,46^, we interrogated whether myelination was affected in the forebrain. To this end, we assessed myelin integrity by performing label-free imaging of compact myelin, using spectral confocal reflectance (SCoRe) imaging of the corpus callosum^47,48^. dLZ/dLZ mice exhibited a 10% decrease in ScoRE-positive area in rostral sections, while a 15% decrease was observed in caudal sections (**Figure 4C-F, Extended Data Fig. 5C**). To determine whether loss of myelin integrity was accompanied by a loss of myelin content, we performed fluoromyelin staining to label myelin lipids directly (**Figure 4G**), which showed a 50% reduction in the corpus callosum. Together, these findings suggest that the animal model recapitulates the reduction in myelin observed in human patients.

**Figure 4:**
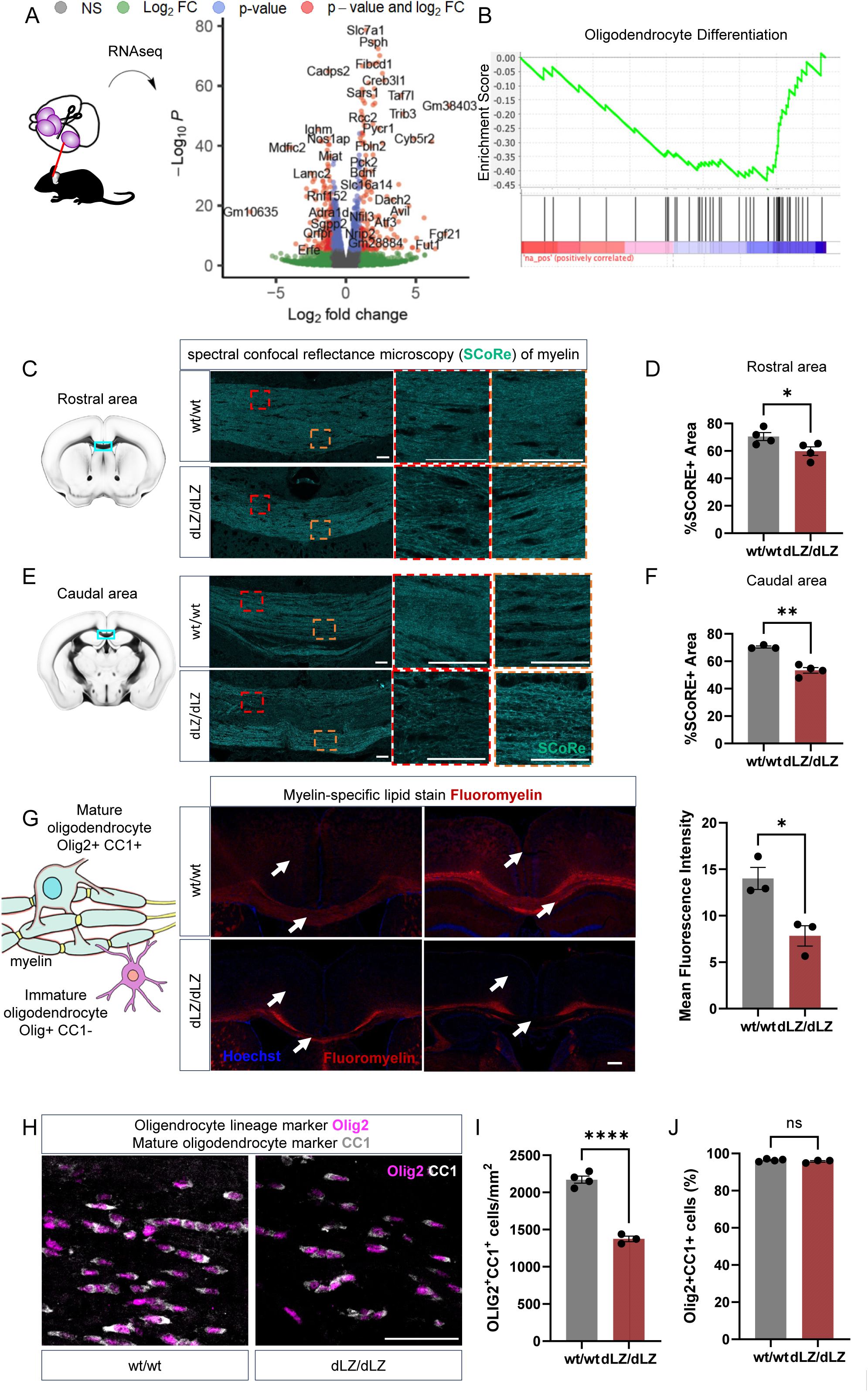
Defective myelination in mice lacking full-length RARS1. A) Volcano plot showing differentially expressed genes in the forebrain as identified by RNAseq. pCutoff=1e-05, Fc cutoff=1. n=3, 3. B) Pathway analysis using ranked differentially expressed genes suggested decreased oligodendrocyte differentiation. C, D) Representative spectral confocal reflectance (SCoRe) images to visualize myelin depositions and integrity. Cryo-embedded corpus callosum of wt/wt and dLZ/dLZ mice in C) rostral sections and D) caudal sections. Bar = 50 µm. Coronal sections illustrations adapted from the Allen Brain Atlas. E) Percentage area positive for SCoRe signal in the corpus callosum of wt/wt and dLZ/dLZ mice. n = 4, 4. Unpaired, two-tailed t test, *p = 0.0471. Insets are magnified images of the squared regions. F) Percentage area positive for SCoRe signal in the corpus callosum of wt/wt and dLZ/dLZ mice. n = 3, 4. Unpaired, two-tailed t test, *p = 0.0011. G) Fluoromyelin staining of myelin in cryosections. F) Mean fluorescence intensity of fluoromyelin signal in the corpus callosum. Bar = 200 µm. n = 3, 3. Unpaired, two-tailed t test, *p = 0.0186. H) Images of cells expressing the oligodendrocyte lineage marker Olig2 (magenta, Oligodendrocyte transcription factor 2) and the marker of mature oligodendrocytes CC1 (white, Quaking 7) in wt/wt and dLZ/dLZ mice. Bar = 50 µm. (I) Number of mature oligodendrocyte cells (Olig2+CC1+) /mm^2^ in wt/wt and dLZ/dLZ mice. n = 4, 3. Unpaired, two-tailed t test, ****p < 0.0001. J) Percentage of mature oligodendrocyte-lineage cells (Olig2+, CC1+ per Olig2+). n = 4, 3. Unpaired, two-tailed t-test, p>0.05. A-J) wt/wt: wildtype RARS1, *Emx1-*Cre+ x Rars1^wt/wt^. dLZ/dLZ: RARS1 delta leucine zipper, *Emx1-*Cre+ x Rars1^loxP/loxP^. Replicates are individual animals. All experiments were performed in 2 months old male mice. C, D, G, H: A representative of at least 3 replicates is shown.

To identify whether the reduction in myelin content is due to changes in oligodendrocyte lineage cells, we quantified cells of the entire lineage (Olig2^+^), as well as those that are mature (Olig2^+^CC1^+^) and immature (Olig2^+^CC1^-^) in the corpus callosum (**Figure 4H, Extended Data Fig. 5D**). dLZ/dLZ mice exhibited a 37% decrease in mature oligodendrocytes (**Figure 4I**). The same proportion of mature CC1+ oligodendrocytes in both wt/wt and dLZ/dLZ suggests that oligodendrocyte maturation was not affected by RARS1 truncation (**Figure 4J**). Staining of Bcas1, a marker for newly formed, myelinating oligodendrocytes, mostly traced patterns comparable to myelination and was decreased (**Extended Data Fig. 5E**). Collectively, these data support that *Emx1*-Cre x Rars1^loxP/loxP^ mice can be used to study hypomyelination.

### Disrupted commitment of neural stem cells to the oligodendrocyte lineage

The significantly reduced oligodendrocyte-lineage population in dLZ/dLZ animals suggested disruptions in earlier neural precursor fate determination, consequently leading to hypomyelination. To further elucidate the cellular origins of the myelin pathology, we cultured cortical precursors isolated at E14.5 from a systemic Cre driver, *Sox2*-Cre^33^ x RARS1^loxP/loxP^ animals, which are systemically expressing dLZ RARS1 only (**Figure 5A**). Animals with systemic RARS1 truncation were used to ensure all cells express only the shorter form of RARS1. Cells were differentiated for one week to recapitulate the *in vivo* sequential generation of neurons and glial cells^49^, allowing us to interrogate late embryonic/neonatal timepoints. Consistent with our findings in adult *Emx1*-Cre x Rars1^loxP/loxP^ mice, oligodendrocyte-lineage cell numbers were strongly reduced in cells derived from dLZ/dLZ animals compared to wt/wt (**Figure 5B**). In turn, proliferating (Ki67-positive) cells decreased, indicating that precursors and/or glial cells were impaired in their proliferation (**Figure 5C**). Antibodies against NeuN, which mark the RNA binding protein Rbfox3, and transcription factor SatB2 were used to identify differences in neuronal cell differentiation (**Figure 5D, E**). Neuronal differentiation was not significantly affected. Altogether, these data highlight early developmental disruptions specifically in the oligodendrocyte lineage caused by RARS1 truncation.

**Figure 5:**
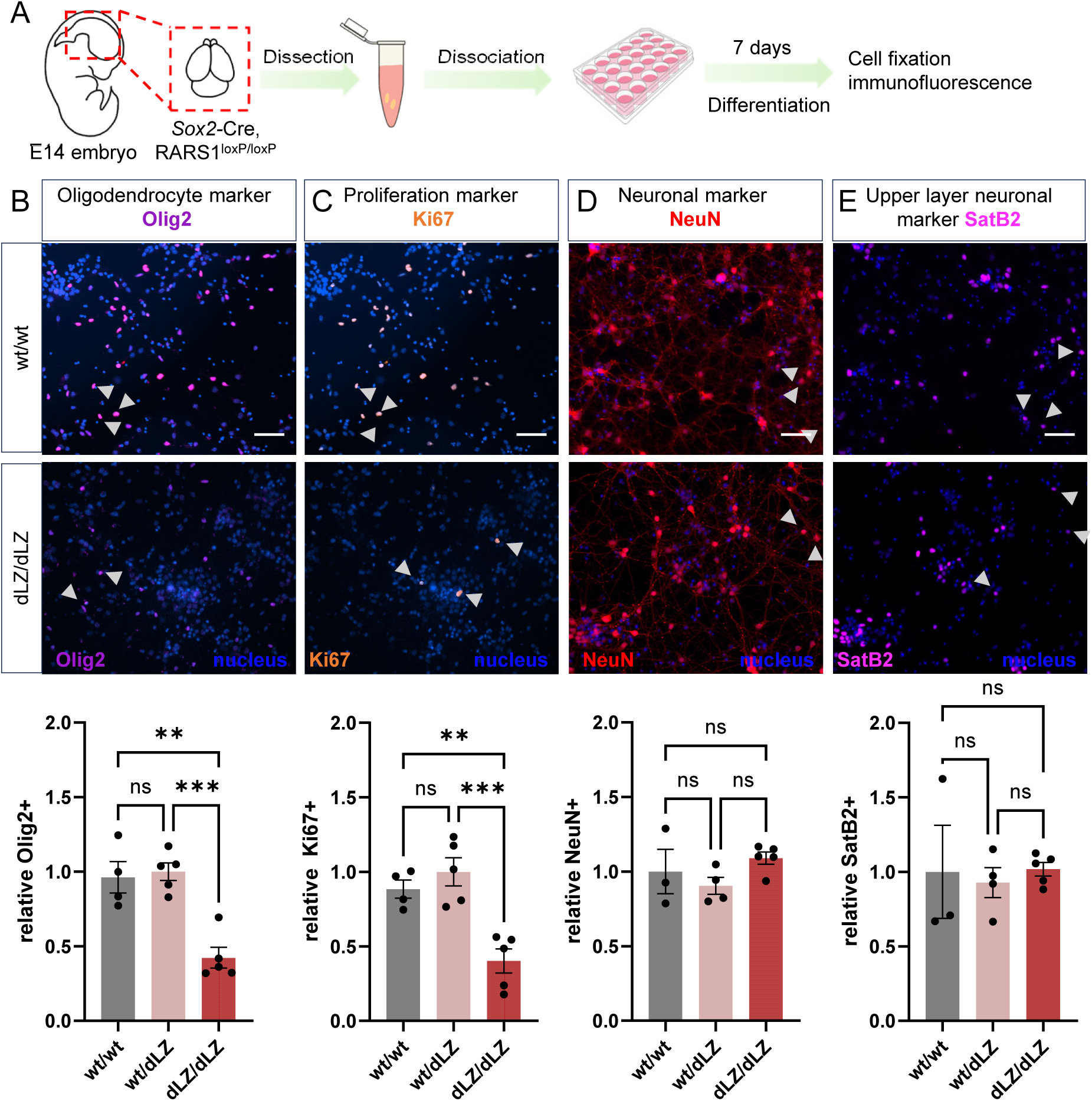
Exclusion of RARS1 from the multisynthetase complex depletes oligodendrocyte-lineage cells during embryonic development. A) Scheme of isolation and differentiation of primary cortical progenitor cells into oligodendrocytes and neurons. (B-E) Upper panels: Representative images of isolated, cultured, and differentiated E14.5 cortical precursors. Hoechst was used to quantify total cell numbers. Bar = 50 μm. Lower panels: B) Relative numbers of Olig2 (Oligodendrocyte transcription factor 2)-positive cells, denoting cells of the oligodendrocyte cell lineage. n=4, 5, 5. Ordinary, one-way ANOVA, multiple comparisons, **p=0.0013, ***p=0.0005. C) Relative number of proliferating cells marked by Ki67 (Proliferation marker protein Ki-67). n=4, 5, 5. Ordinary, one-way ANOVA, multiple comparisons, **p=0.0054, ***p=0.0007. D) Relative numbers of NeuN (RNA binding protein fox-1 homolog 3)-positive cells, denoting cells of the neuronal lineage. n=4, 5, 5. Ordinary, one-way ANOVA, multiple comparisons, p>0.05. E) Relative numbers of SatB2 (DNA-binding protein SATB2)-positive cells, denoting cells of the neuronal lineage, specifically of the upper layer. n=4, 5, 5. Ordinary, one-way ANOVA, multiple comparisons, p>0.05. A-E) wt/wt: wt RARS1, systemic inclusion of RARS1 exon 2. dLZ/dLZ: RARS1 delta leucine zipper, systemic exclusion of RARS1 exon 2. Mice originated from *Sox2-*Cre x Rars1^loxP/loxP^ and were bred as wt/dLZ animals following F1. Replicates are individual embryos. Sex was determined but results were not stratified by sex.

### Altered subcellular localization of N-terminally truncated RARS1

Our previous work proposed that the MSC plays a role in regulating the subcellular distribution of ARSs^21^. Hence, we were interested in studying the consequences of RARS1 exclusion from the MSC on RARS1’s subcellular localization. We again made use of cultured, differentiated E14.5 cortical precursors as a model for different primary cell types. We co-stained cells with antibodies against RARS1 and the ribosomal protein S6 to mark ribosomes. Of note, RARS1 intensity was higher in neurons than in other cell types (**Extended Data Fig. 6A**), which made quantitative analysis in those cells challenging. We therefore assumed that dysregulation in subcellular localization due to a loss of RARS1 from the MSC would be comparable between cell types. We employed Nissl staining, which binds to negatively charged biomolecules in cells such as DNA and RNA, to trace the overall cell shape. In wt/wt cells, both RARS1 and S6 were found mostly in the cell body, and Manders coefficients indicated a high degree of colocalization (67%), suggesting that the translational components clustered together (**Figure 6A, B, Extended Data Fig. 6B**). However, in dLZ/dLZ cells, RARS1 and S6 differed significantly in their colocalization, as evidenced by a significantly reduced Manders coefficient. RARS1 distribution within dLZ/dLZ cells was less organized and increased nuclear localization observed. As different cell types intermingled in cellular projections, we refrained from quantifying colocalization. In summary, a difference in subcellular localization was clearly visible when comparing wt/wt and dLZ/dLZ RARS1 in cell bodies.

**Figure 6:**
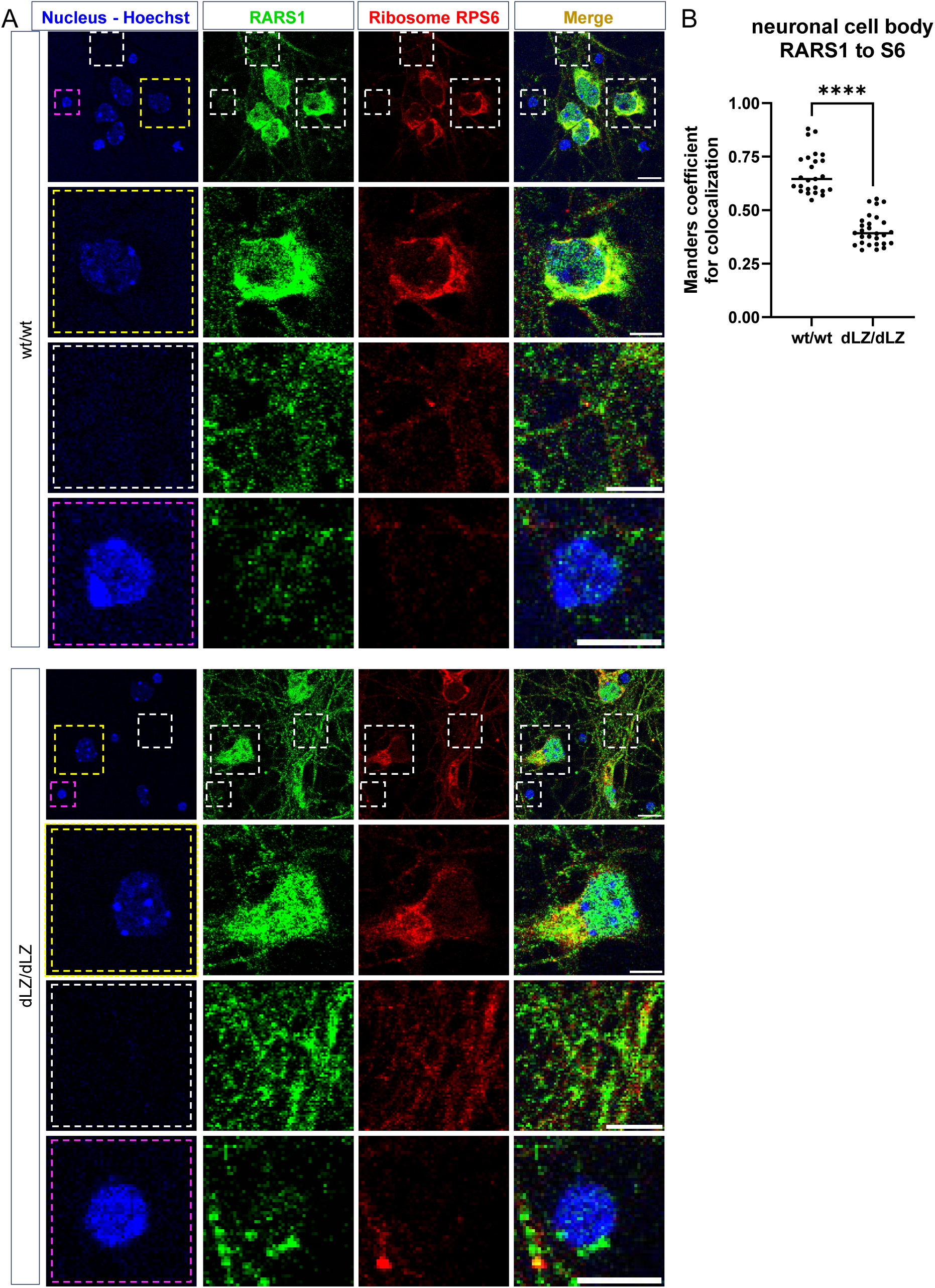
Altered subcellular localization of RARS1 following its N-terminal truncation. A) Representative images of isolated, cultured, and differentiated E14.5 cortical precursors. RARS1 and S6 were visualized using immunofluorescence. Hoechst was used to identify nuclei, indicating cell bodies. Yellow frame: Neuronal cell bodies. White frame: neuronal cell projections. Pink frame: Other cell bodies, including oligodendrocytes. Bar = 10 μm in the overview image, 5 μm in the zoomed in image. B) Manders coefficient indicating colocalization of RARS1 relative to S6 in neuronal cell bodies. n=25, 28. Unpaired, two-tailed t test, ****p<0.0001. A, B) wt/wt: wt RARS1, systemic inclusion of RARS1 exon 2. dLZ/dLZ: RARS1 delta leucine zipper, systemic exclusion of RARS1 exon 2. Mice originated from *Sox2-*Cre x Rars1^loxP/loxP^ and were bred as wt/dLZ animals following F1. Replicates are individual embryos. Sex was determined but results were not stratified by sex.

### Exclusion of RARS1 from the multi-tRNA synthetase complex induces integrated stress response signaling

Given the disruption of RARS1 subcellular organization, we were interested in further understanding molecular consequences. Upregulated transcripts identified by RNAseq were also enriched in those associated with cellular stress response, specifically ER stress pathways, suggesting that protein synthesis of secreted and membrane proteins could be perturbed (**Figure 7A, Extended Data Fig. 7A**). We therefore investigated the integrated stress response (ISR) pathway, as it responds to ER stress and is commonly dysregulated in disorders caused by ARS mutations, as they can cause a lack of charged tRNAs^50,51^. Oligodendrocytes, which were most affected in our mouse model, are particularly sensitive to ISR^52^. The stress-induced transcription factor ATF4 was upregulated on mRNA level (**Supplementary Table 4**) and immunoblot in forebrain lysates confirmed that ATF4 protein levels were increased (**Figure 7B**). Concomitantly, we observed a concerted upregulation of ATF4 target genes, highlighting a sustained integrated stress response in adult mice (**Figure 7C**).

**Figure 7:**
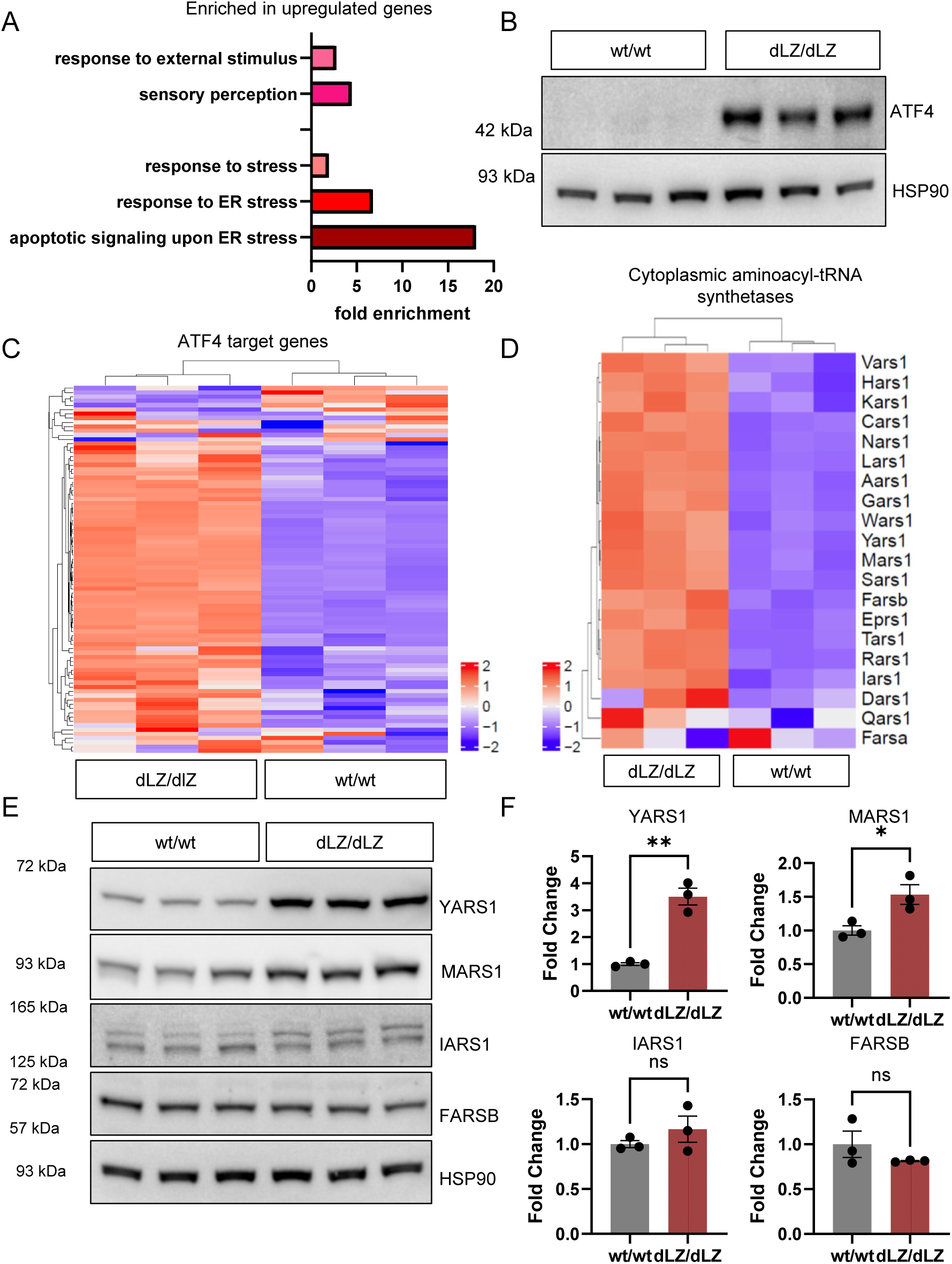
Exclusion of RARS1 from the multisynthetase complex activates cellular stress response. A) Pathway enrichment using differentially upregulated genes found in transcriptome analysis using RNAseq (at least 1.5-fold change, padj<0.05) showed an induction of cellular stress response, specifically ER stress signaling. B) Immunoblot of ATF4 in forebrain lysates. HSP90 was used as a loading control. C) Z-scaled heatmap of ATF4 target gene mRNA levels. D) Z-scaled heatmap of cytosolic aminoacyl-tRNA synthetases mRNA levels. E, F) Immunoblot and quantification of select aminoacyl-tRNA synthetases in 2-month-old male mice forebrain lysates. E) Immunoblot, HSP90 was used as a loading control. F) Densitometric quantification of aminoacyl-tRNA synthetases immunoblots. All values were normalized to HSP90 levels. Unpaired, two-tailed t-tests, YARS1: **p = 0.0015, MARS1: *p = 0.0308, IARS1: p > 0.05, FARSB: p > 0.05. A-F) wt/wt: wildtype RARS1, *Emx1-*Cre+ x Rars1^wt/wt^. dLZ/dLZ: RARS1 delta leucine zipper, *Emx1-*Cre+ x Rars1^loxP/loxP^. Replicates are individual animals. All experiments were performed in 2 months old male mice.

Cytosolic aminoacyl tRNA synthetases are targets of the ATF4/CHOP pathway^53^. Strikingly, mRNA levels of almost all cytosolic ARS were upregulated, while their mitochondrial counterparts were not uniformly affected (**Figure 7D, Extended Data Fig. 7B**). To assess whether these expression changes led to altered protein levels, we performed immunoblot against multisynthetase complex-bound and free-standing ARS. We found that levels of the canonical ATF4 effector YARS1 were significantly increased, while other ARS changes (MARS1, IARS1, FARSB) were more subtle (**Figure 7E, F**). Of note, IARS1 showed isoform specific changes, which may relate to MSC integrity^54^. In addition to ARS, nearly all ribosomal subunits also displayed upregulation on mRNA level in dLZ/dLZ animals (**Extended Data Fig. 7C, D**), suggesting the wide-spread dysregulation of cellular homeostasis genes. Of note, this likely explains the increase of RARS1 activity, protein, and mRNA observed in dLZ/dLZ animals (**Figure 1F-H**), as RARS1 was among the affected ARS genes.

As RARS1 has been associated with alternative mRNA splicing^55^, we assessed differential changes in splicing events in dLZ/dLZ vs wt/wt mice. Genes with changes in mRNA splicing were enriched in those involved in neuronal signaling and development, mRNA localization and splicing regulation, as well as synapse function (**Extended Data Fig. 7E**). However, splicing changes were minor overall and the differences in ‘percent spliced in’ (PSI) between wt/wt and dLZ/dLZ were < 10% for all event types (**Extended Data Fig. 7F, Supplementary Table 5**). These somewhat subtle differences could be reflective of insufficient coverage due to the heterogenous RNA input or that overall changes were predominantly affecting mRNA expression as opposed to processing.

Together, this suggested that mRNAs associated with cell stress and protein synthesis were regulated in dLZ/dLZ animals, which could cause or indicate a change in cellular homeostasis upon exclusion of RARS1 from its protein complex.

### Exclusion of RARS1 from the multisynthetase complex disrupts local protein synthesis at cell projections

Given that bulk RARS1 enzymatic activity was increased (**Figure 1F**), we were surprised by the consistent indication that cellular stress pathways, commonly associated with a lack of charged tRNAs, were upregulated (**Figure 7B, C**). We therefore aimed to probe newly synthesized proteins to assess whether translation would be affected. We made use of a recently developed bioconjugation reaction^56^, which allows for the attachment of a strained alkyne to an unnatural amino acid, a tellurophene-based phenylalanine analogue (TePhe). Reaction with the strained alkyne allowed for the installation of fluorescent moieties, such as TAMRA, or biotin-handles for enrichment, a procedure we named TeACAT: <u>Te</u>llurophene-<u>A</u>lkyne <u>C</u>ycloaddition-mediated <u>A</u>mino acid <u>T</u>agging^57^. TeACAT benefits from the similarity of TePhe to Phe, which allows its incorporation into proteins without the need for concomitant depletion of the canonical amino acid or genetic manipulation. We injected TePhe twice over the course of 8 h (**Figure 8A**) and quantified total TePhe incorporation using TAMRA-labeling and fluorescence (**Extended Data Fig. 8A**). Overall protein synthesis was comparable, suggesting that bulk translation was not affected. Bulk protein synthesis quantification in lysates was confirmed by applying TeACAT to tissue sections, which showed overall uniform and comparable protein synthesis when assessing whole mount brain sections (**Extended Data Fig. 8B, C**). We then probed for the identity of newly synthesized proteins using the same animals and pulled down TeACAT proteins from the forebrain by attaching a biotin handle (**Figure 8A**). We were able to identify ∼2000 proteins, 74 of which were significantly changed when comparing dLZ/dLZ to wt/wt proteomes (>1.5 fold change, *p*-value <0.05, **Supplementary Table 6-9**). Interestingly, we found an enrichment for proteins associated with cell projections and other compartments of the cell, which are known to rely on local translation (synapse, plasma membrane) (**Figure 8B**). By comparing changes on mRNA level with those found in newly synthesized proteins, we found that mRNA levels of these differentially synthesized proteins remained unchanged (**Figure 8C**). This strongly suggests that while bulk translation was not affected, local translation at cellular projections might be downregulated in RARS1 dLZ/dLZ forebrains.

**Figure 8:**
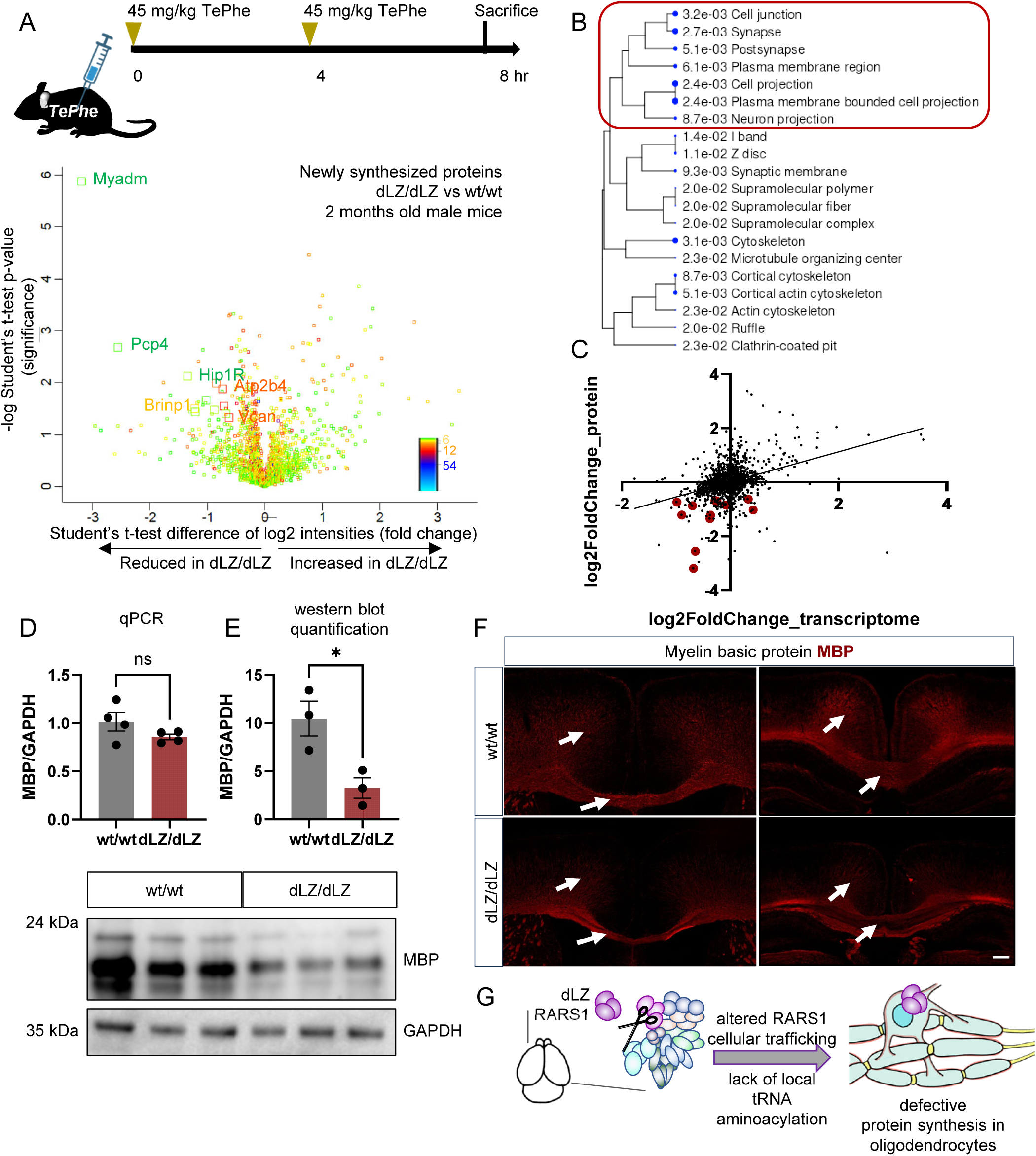
Exclusion of RARS1 from the multisynthetase alters protein synthesis of locally translated neural gene transcripts. A) Upper panel: Administration regime of TePhe to mark newly synthesized proteins. Intraperitoneal injections were performed at 0 and 4 h time points. Tissue collection was performed 8 hours after the first injection. Lower panel: Volcano plot representing newly synthesized proteins enriched by pulldown of the unnatural amino acid TePhe incorporated into proteins. n=4, 3. B) Enrichment analysis to identify cell compartments assigned to TePhe labeled proteins. C) Comparison between RNAseq transcriptome and TeACAT proteome of newly synthesized proteins. Genes/proteins labeled in red correspond to genes associated with cell projections. Rsquare: 0.147. Sy.x: 0.4620. Slope: 0.5566. D) qPCR analysis of MBP mRNA levels showed unchanged MBP expression. GAPDH was used as a control. n=4, 4. Unpaired, two-tailed t test, p>0.05. E) Immunoblot of myelin binding protein (MBP) and densitometric analysis showed a reduction in dLZ/dLZ animals compared to wt/wt. GAPDH was used as a loading control. n=3, 3. Unpaired, two-tailed t test, *p = 0.0263. F) Immunofluorescence of Myelin basic protein (MBP) in forebrain cryosections. The solid arrow indicates the cortex and the dashed arrow indicates the corpus callosum. Hoechst was used as a counterstain to mark nuclei. Bar = 200 µm. A representative of at least 3 experiments is shown G) Scheme: Exclusion of RARS1 leads to altered cellular localization and affects local protein synthesis in oligodendrocyte cellular projections, leading to a reduction in oligodendrocyte lineage cells and ultimately white matter brain disorder symptoms. A-F) wt/wt: wildtype RARS1, *Emx1-*Cre+ x Rars1^wt/wt^. dLZ/dLZ: RARS1 delta leucine zipper, *Emx1-*Cre+ x Rars1^loxP/loxP^. Replicates are individual animals. All experiments were performed in 2 months old male mice.

We further interrogated whether protein synthesis was especially dysregulated in specific cell types: proteins with reduced expression were associated with oligodendrocytes (**Extended Data Fig. 8D**). In line with RNAseq results, the synthesis of most aminoacyl-tRNA synthetases was increased (**Extended Data Fig. 8E**). This points towards the disruption of localized protein synthesis in dLZ/dLZ animals. Looking back into the RARS1 interactome, we noted additional RARS1 interaction partners in dLZ/dLZ lysates: In addition to a loss of MSC-bound ARS as interactors, increasing interaction with free-standing ARS was observed (YARS1, WARS1, AARS1, and NARS1, **Extended Data Fig. 8F-H).** An increasing number of proteins with nuclear localization were found in addition to cytosolic and myelin sheath proteins, in line with microscopy images obtained from primary cells (**Figure 6**, **Extended Data Fig. 8F-H)**. Together, this enforces a change in RARS1 localization upon its exclusion from the MSC.

We decided to investigate one exemplary protein in the depth: Myelin basic protein (MBP) is locally translated after transport of its translationally inhibited mRNA to axon-glial contact sites and is critical for myelination^58^. qPCR revealed that MBP mRNA levels were unchanged (**Figure 8D**), however, immunoblot quantification of MBP in forebrain lysates showed a reduction on protein level to less than 50% of wt/wt animals (**Figure 8E**). Immunofluorescence staining of MBP illustrated hypointense cortical and corpus callosum signal (**Figure 8F, Extended Data Fig. 8I**), which aligned with the decompaction of myelinated tracts observed with SCoRe imaging (**Figure 4A**). These data suggest an impairment in MBP protein synthesis, possibly due to a reduction of local RARS1 availability, leading to a reduction of brain white matter (**Figure 8G**). This directly links a molecular mechanism, ARS complex formation and subsequent cellular localization, to disease phenotypes.

## Discussion

Here we investigated the importance of the multi-tRNA synthetase complex for neurodevelopment and found that if complex formation is disrupted, brain white matter decreased concomitantly to a reduction in mature oligodendrocytes. At the molecular level, RARS1 localization within the cell was perturbed, suggesting a role of the MSC in RARS1 subcellular trafficking. In consequence, protein synthesis in cellular projections was altered, ultimately causing hypomyelination.

The RARS1 dLZ/dLZ mouse model mimicked phenotypes observed in human patients, suggesting its suitability as a preclinical model: the reduced brain size observed in dLZ/dLZ animals was consistent with microcephaly reported in patients suffering from loss of full-length RARS1^28,59^. Both male and female dLZ/dLZ mice exhibited gross motor coordination issues as revealed by behavioral assays, including difficulty balancing on a ledge (**Figure 2B**) and worsened rotarod performance in male mice (**Figure 2C**). Observed anatomical changes reflected on the role of distinct brain regions on controlling behavior: a reduction in cortical thickness, especially in cortical regions important for motor control, have been associated with motor deficits including reduced dexterity in human patients^60^. These effects are most likely intertwined with hypomyelination-related effects such as finer sensorimotor deficits in a hypomyelination model and impaired motor behavior in demyelination models^61,62^. Unexpectedly, dLZ/dLZ animals exhibited higher velocity than their wild-type counterparts in open field testing which may be attributed to the increased anxiety-like behavior in these animals (**Figure 2D, 2E**). Anxiety-like behavior has been previously associated with cortical thinning and microcephaly^63,64^. Altered neural circuitry may also play a role in the observed behavioral defects, as *Emx1*-controlled RARS1 truncation targets excitatory neurons but not inhibitory neurons^65^ which may lead to excitatory-inhibitory imbalances in dLZ/dLZ animals.

In contrast to loss-of catalytic function variants in several aminoacyl-tRNA synthetase-related disorders^66^, truncation of RARS1 in our model increased bulk aminoacylation activity of RARS1 (**Figure 1F**) despite striking changes in brain anatomy between wt/wt and dLZ/dLZ mice (**Figure 3A, B**). This suggested that reduced overall enzymatic activity was not causal for the observed phenotype. Our results echo clinical findings where loss of full-length RARS1 in patients suffering from hypomyelination did not affect bulk catalytic activity in patient fibroblasts despite severe clinical presentation^28^.

Measurements in bulk lysates may not reflect subcellular changes in RARS1 availability, however. We found that upon exclusion from the multisynthetase complex, RARS1 accumulated in neuronal nuclei, showing clearly altered subcellular trafficking compared to wt/wt neurons (**Figure 6A**). With altered cellular distribution of RARS1, the availability of RARS1-charged tRNAs for translation of distal mRNAs may be lower, thereby impairing translation in cell projections. Indeed, tagging of newly synthesized proteins using TeACAT suggested that production of proteins localized to cell projections was reduced (**Figure 8A-C**). This is especially notable as neurons rely heavily on localized translation for protein synthesis due to their extensive cell projections^67–73^. In addition to neurons, oligodendrocytes are particularly vulnerable to disruptions in local protein synthesis, as proteins associated with myelin formation are synthesized in distal sites of the ER^71,74^.

Our data also sheds light on the importance of RARS1 localization in oligodendrocyte biology. Altered RARS1 localization in our model seemingly perturbed local translation of mRNAs (**Figure 8A-C**), including myelin basic protein: MBP protein levels were decreased while its mRNA levels were unchanged (**Figure 8D-F**). Disturbed localized translation in oligodendrocytes may be further exacerbated by lower RARS1 levels compared to neurons (**Extended Data Fig. 6A**). This might explain why oligodendrocytes were especially vulnerable to local deficits in protein synthesis and could contribute to the reduction in mature oligodendrocyte numbers (**Figure 4 H-J**), ultimately leading to a white matter defect as the predominant phenotype in patients.

This vulnerability is also reflected in early developmental stages, where oligodendrocyte lineage cells in embryonic cultures and adult mice were reduced (**Figure 4, 5B**). In contrast, neuron numbers were mostly unchanged both in embryonic cultures and in adult mice (**Figure 3D-F, 5D, E**). This observation may aid in differentiating the effects of RARS1 truncation in neurons and oligodendrocytes. While oligodendrocyte lineage commitment might have been affected, a loss in myelin compaction and MBP specifically may also reflect disturbed neuron guidance during myelination (**Figure 3A-D**).

Perturbed localized translation could directly explain the upregulation of ER stress pathways and the integrated stress response (**Figure 7A-C**). Accumulation of uncharged tRNAs triggers the integrated stress response and mislocalization of RARS1 could cause a local lack of charged tRNAs. Currently, we are missing direct methods to assess tRNA and tRNA aminoacylation with subcellular resolution, necessitating measurement of newly synthesized proteins as a proxy for local tRNA availability instead (**Figure 8A-C**).

We observed upregulation of most ARS as well as ribosomal components (**Figure 7D-F, Extended Data Fig. 7D**), which could be a systemic compensatory response to the local lack of charged tRNA. ISR activation is in line with previous reports which identified aminoacyl-tRNA synthetases as effectors of stress response^8^. For instance, YARS1 translocates to the nucleus upon stress and regulates the expression of translation-related genes upon prolonged cellular stress^75^. The accumulation of RARS1 in the nucleus and its increased interaction with YARS1 upon truncation could point to mechanisms promoting YARS1-mediated cell survival as well as a rerouting of RARS1 to ex-translational and noncanonical functions, such as the regulation of alternative splicing^55^. Elevated RARS1 levels could also perturb cellular arginine levels, thereby restricting metabolite availability for other signaling pathways^76–79^.

Altogether, we identified the necessity of ARS complex formation, and especially the integration of RARS1 therein, for mammalian neurodevelopment. RARS1 complex integration was a prerequisite for local protein synthesis, and oligodendrocyte lineage cells were especially sensitive to its disruption. Overall, our findings suggest a novel molecular mechanism underlying human disease.

## Material and Methods

### Mouse husbandry

All experimental protocols were approved by the Health Sciences Local Animal Care Committee at the University of Toronto (AUP 20012885). Mice were housed under standard laboratory conditions with access to food and water. The housing room underwent 12h dark/light cycles. Rars1^loxP/loxP^ (registered as Rars1^Tm1Cuiut^) mice were generated by the Salk transgenic animal facility using CRISPR/Cas9. Mice systemically lacking full-length RARS1 were obtained by crossing B6.Cg-Edil3Tg(Sox2-cre)1Amc/J mice with Rars1^loxP/loxP^ and maintained as dLZ/wt crossings. Mice lacking full-length RARS1 in the forebrain were obtained by crossing B6.129S2-*Emx1tm1(cre)Krj*/J (Jackson Laboratories) to Rars1^loxP/loxP^.

### Genotyping

Genomic DNA was extracted from mouse ear snips using DNA Quick Extract^TM^ Solution (Bioresearch Technologies) as per the manufacturer’s instructions. LoxP genotyping was performed by PCR amplifying the genetic loci with loxP site insertions. Amplicon size was used to assess the presence or absence of loxP sites. Cre genotyping was performed by PCR targeting the CreA gene locus.

### Size exclusion chromatography

Brain tissue was homogenized in 10x volume of MSC lysis buffer (20 mM Tris–HCl, pH 7.5, 1 mM EDTA, 150 mM NaCl, 1% NP-40 and Protease Inhibitor Cocktail cOmplete™, Mini, EDTA-free Protease Inhibitor Cocktail, Roche). The lysate was incubated at 4°C for 30 minutes, then centrifuged at 14,000x g for 20 min to remove solid debris. The supernatant was loaded onto a Superdex 200 Increase 10/300 GL Prepacked SEC Column (Cytiva) equilibrated with PBS (Gibco) and 1 mL fractions were collected. 25 μL of each fraction was analyzed by immunoblot. Blots were exposed together during imaging, therefore signal intensities between individual blots are comparable.

### Immunoblot

Brain cortices were snap-frozen in liquid nitrogen immediately after harvesting. Frozen brain tissue samples were homogenized in TBS or RIPA lysis buffer (from 10x: 0.5 M Tris/HCl, pH = 7.4, 1.5 M NaCl, 2.5% deoxycholate, 10% NP-40, 10 mM EDTA) with protease inhibitor cocktail. Lysates were incubated at 4°C for 30 minutes, then centrifuged at 13,000 xg for 20 min. The supernatant was mixed with 5× SDS loading buffer, boiled for 10 min, and 20 ug – 40 μg of protein were loaded on a 4–12% gradient gel. Total protein was quantified by Bradford Assay (BioRad). Protein was transferred to nitrocellulose membranes using an iBlot2. Membranes were blocked in 1-5% milk/TBST for 1 hour and incubated in the respective antibody diluted in 5% BSA/TBST overnight at 4°C. The following antibodies were used: RARS1 (Biorbyt, 1:1000), MBP (Abcam, 1:2000), IARS1 (Proteintech, 1:2000), MARS1 (Proteintech, 1:3000), QARS1 (Proteintech, 1:1000), QARS1 (ThermoFischer, 1:1000), YARS1 (SCBT 1:5000), TARS3 (Proteintech, 1:1000), FARSB (Proteintech, 1:2000), IARS1 (Proteintech, 1:1000), ATF4 CST, 1:1000), HSP90 (CST, 1:1000), GAPDH (CST, 1:5000). The next day, membranes were washed 3× in TBST for at least 10 min and incubated for at least 90 min in secondary antibody diluted in 1% milk/TBST (rabbit secondary: 1:5000, mouse secondary 1:10,000, both Invitrogen). Membranes were washed again 3× for at least 10 min. Visualization was done using ECL substrate (Cell Signaling) on a BioRad Imager. Immunoblots were exposed just below saturation to take advantage of the maximum dynamic range. Signal intensities between individual blots are therefore not comparable.

### qPCR

cDNA was synthesized from 1 µg of total RNA using the RevertAid First Strand cDNA Synthesis Kit (Thermo Scientific) with oligo(dT) primers, following the manufacturer’s instructions. Quantitative PCR (qPCR) was performed on a LightCycler® 480 System (Roche) in 25 µL reactions containing 12.5 µL of 2× PowerUp SYBR Green Master Mix (Applied Biosystems), 500 nM of each primer, and 50 ng of cDNA with nuclease-free water added to volume. Cycling conditions were 95 °C for 10 min, followed by 50 cycles of 95 °C for 15 s, 60 °C for 30 s, and 72 °C for 30 s. All reactions were run in technical triplicates with n≥4 biological replicates. Gene expression was normalized to GAPDH, and relative fold changes were calculated using the 2^ΔΔCt^ method.

### Aminoacylation assays

RARS1 tRNA charging activity was measured using a filter-binding assay. Mouse cortices were dissected and homogenized as described above in MSC lysis buffer supplemented with protease inhibitors. The enzymatic reactions were started by adding 5 µL of lysate to a mixture of in vitro transcribed human arginyl-tRNA, 100 mM HEPES pH 7.5, 20 mM KCl, 2 mM DTT, 10 mM MgCl_2_, 0.004 mg/mL pyrophosphatase (NEB), 20 µM cold L-arg, 5 µM ^3^H-L-arg, 4 mM ATP. The reactions were incubated at 37 °C and quenched by precipitating the tRNAs with 20% cold TCA in a 96-well filter plate after 1 min, 5 mins and 15 mins. The filters were washed four times in ice cold 100 mM L-arg, 5% TCA, then washed once with 95% ethanol. Filters were dried, then charged tRNAs were hydrolyzed by adding 80 µL of 0.1 M NaOH. The released radiolabeled arginine was collected in scintillation liquid. After mixing, the plate was read by a liquid scintillation counter.

### Tissue Collection and Processing

For immunofluorescence and SCoRE imaging, mice were anesthetized using isoflurane, then intracardially perfused with 20 mL of PBS followed by cold 4% paraformaldehyde/PBS solution. Brains were harvested and post-fixed overnight at 4 °C in 4% PFA before being transferred in 30% sucrose for approximately 72 hours. Brains were embedded in OCT medium, then frozen on dry ice. 18 µm coronal brain sections were obtained and stored at -80 C until further processing.

### Luxol Fast Blue and H&E staining

Mice were euthanized by CO_2_ exposure. Brains were harvested and directly drop-fixed in 4% PFA for 60-72 hr with one change of fixative halfway through the fixation period. Brains were rinsed in PBS, followed by division of the forebrain into 3 roughly equal-thickness coronal regions using a brain matrix, then dehydrated and embedded according to published procedures^80^. Tissues were sectioned into 4 µm slices for staining with Luxol Fast Blue (Thermo Scientific), which followed standard procedures: after incubation at 60 °C for 30 minutes, brain sections were deparaffinized and rehydrated then submerged in 0.1% Luxol Fast Blue solution for 16 hours at 56 °C. Sections were rinsed in 95% ethanol, then differentiated in 0.05% lithium carbonate before mounting in ProLong Gold mounting medium (Invitrogen). For H&E staining, 5µm thick sections were baked at 60 °C for 30 minutes, then deparrafinized in an ethanol series. Sections were stained with hematoxylin for 4 mins, then blued in Scott’s tap water substitute (20 g/L MgSO_4,_ 3.5 g/L NaHCO_3_ in water, H from Bioshop) for 30 s, then stained with Eosin Y (2.5 g/L in 80% v/v reagent alcohol, 1% v/v acetic acid in water, E from Bioshop) for 3 mins. Sections were washed in 95% ethanol, then dehydrated in 100% ethanol, then rinsed three times in xylene before mounting in resinous DPX medium (Sigma).

### Behavior Testing

Behavioral tests were performed on littermates. Mice were allowed to acclimate to the testing environment for at least 10 minutes prior to testing. Tests were performed at a similar time of the day between cohorts.

### Rotarod

Mice were placed on a rotating rod (Data Sciences International) accelerating from 4 rpm to 60 rpm over 60 seconds. Their latency to fall was recorded. Measurements were done in duplicates spaced out by at least 20 minutes.

### Grip strength

Mice were allowed to grasp a metal grid attached to a horizontal grip strength meter (Data Sciences International). They were subsequently pulled backwards in the horizontal plane until they released their grasp. Maximum grip strength was measured as the peak force applied to the grid just before the release. The measurement was performed five consecutive times, and the average of the five measurements was reported.

### Ledge navigation

Mice were placed on a cage ledge and left to walk along the ledge. They were assigned a score based on their ability to navigate along the ledge. Mice who walked without losing their balance and who lowered themselves gracefully using their paws were assigned a score of 0. Incrementally higher scores were given based on the severity of the walking defects ranging from exhibiting lack of balance, over-reliance on their head as opposed to hind limbs for lowering themselves into the cage, inability to move in the most severe cases. Videos and scores were reviewed by a second experimentalist.

### Gait Analysis

Mice front and hind paws were painted with contrasting inks (Wordsworth), then released on a straight 7 cm wide tunnel lined with grid paper (Geyer) and long enough for the mice to take at least 5 steps. Footprint analysis was done on the markings: stride length was obtained by measuring the distance between two sequential footprints created by the same foot. Measurements were averaged over 5 gait cycles.

### Open field testing (OFT)

Mice were tested in a square open-field arena whose floor was marked as a 3×3 macroscopic grid; each square was further subdivided into a 4×4 grid, yielding a 12×12 fine grid with each cell measuring 1 inch (2.54 cm) per side. Behavior was recorded from overhead (∼30 fps) under uniform illumination. Videos were analyzed in DeepLabCut ^45^ (ResNet-50), trained on 20 manually labeled frames per video to track the nose and tail base. Position was taken as the body centroid (mean of nose and tail-base coordinates). Arena bounds were defined from the grid; pixel-to-length calibration was 84 px = 1 inch (0.030238 cm·px⁻¹). Occupancy was computed on the 12×12 lattice, with the “perimeter” (thigmotaxis zone) defined as the outermost two fine-grid cells along each wall; remaining cells comprised the center, with corners analyzed separately. Outcomes included time in perimeter, center, and corners; center-entry counts; total distance traveled; average speed. To accommodate variable video lengths, metrics were reported both in seconds and as fractions of tracked time, with counts/distances expressed per second.

### Immunofluorescence

Cryosections were dried for 30 min at 37°C then rehydrated in PBS for 5 min. Sections were then blocked in 5% goat serum and 0.3% Triton X-100 in PBS for 1 h. NeuN (EMD Millipore, 1:200) was applied overnight at 4°C in a humified chamber. For myelin proteins MBP (Abcam, 1:500) and CNP staining (Sigma, 1:1000), cryosections were submerged in methanol at –20°C for 10 mins prior to blocking as described above, and the antibodies were applied overnight at room temperature. For Olig2 (R&D systems, 1:100), APC clone CC1 (EMD Millipore,1:300) and BCAS1 (SYSY, 1:100) staining, sections were air dried for 15 mins, then heat-antigen retrieval was performed by boiling the sections for 5 min in 10 mM sodium citrate pH 5 buffer. Blocking was performed in 5% normal horse serum + 0.3% Triton X-100 in TBS before the antibodies were applied overnight at room temperature. The next day, sections were washed 3 times in PBS-T (PBS + 0.1% Triton X-100) or TBS-T (TBS + 0.1% Tween-20) before secondary antibodies were applied in blocking solution for 2 h at room temperature. Sections were washed 3 times in PBS or TBS before counterstaining with Hoechst 33258 for 5 min, washed twice in PBS, and mounted in ProLong Gold (Invitrogen) mounting medium. Antibodies are listed in **Supplementary Table 9**.

### Spectral Confocal Reflectance Microscopy (SCoRE)

All reflectance images of myelin were captured on a Leica Stellaris 8 Falcon confocal microscope with an acoustic-optical beam splitter (AOBS) with an 85/15 reflectance/transmission ratio. The AOBS collected data from laser wavelengths of 488 nm, 568 nm, and 633 nm using three HyD photodetectors with a very narrow detection bandwidths of 5 nm centered around the chosen laser wavelengths. Detection bands were set to 485–490 nm, 565–570 nm, and 630–365 nm for each laser, respectively. Laser and gain settings were initially set at their minimum, and then slowly increased until the first over-saturated pixel appeared. Then, the gain was slightly lowered until the first disappearance of the over-saturated pixel. Microscope settings were kept the same across images of the same cohort. For this, laser/gain settings were set for the first image and kept the same for all subsequent images. A 40X oil immersion objective (NA1.3) was used. Nuclei staining with DAPI was imaged by sequential scanning using conventional fluorescence acquisition in a separate sequence capturing fluorescence light.

### SCoRE Image Analysis

Images from the 488 nm, 568 nm, and 633 nm channels were merged. The composite image was analyzed by setting a threshold for pixels positive for myelin. The percentage of pixels above the threshold were quantified as the percent area positive for myelin.

### Fluoromyelin Staining

Cryosections were dried for 30 min at 37°C then rehydrated in PBST for 20 min. 1:200 Fluoromyelin Red (ThermoFisher, 1:200 in PBS) was applied for 20 mins. Sections were washed three times in PBS, then counterstained with Hoechst 33258 for 5 min, washed twice in PBS, and mounted in ProLong Gold mounting medium (Invitrogen).

### Antibody expression and purification

DNA sequences encoding anti-RARS1 single chain variable fragments (scFv) were obtained from ^81^. scFv sequences were cloned into plasmid backbones containing antibody mouse IgG light and heavy chains. Anti-RARS1 antibodies were expressed in HEK293T cells. 5 hours before transfection, media was exchanged to 25 μM chloroquine diphosphate (Fisher Scientific) in DMEM. For expression in a 10 cm dish, diluted light chain and heavy chain plasmids (18.9 µg total) were mixed dropwise with diluted polyethylenimine (PEI, linear, average Mn 15,000, 1 mg/mL, Sigma Aldrich) in Opti-MEM at a ratio of 1:3 DNA:PEI and incubated for 15 minutes at room temperature before addition to cells. 18 hours after transfection, media was replaced with serum free media. Cells were incubated for 4 days to condition media. For antibody purification, protein G bead slurry (GenScript) was pre-equilibrated in 10 mM HEPES pH 7 and conditioned media was added. After one hour of gentle agitation at 4 °C, beads were washed in 10 mM HEPES pH 7 and antibody was eluted in 0.1 M glycine/HCl, pH 3.0, through a 0.22 µm spin filter into neutralization buffer (1 M Hepes, pH 9). Purified anti-RARS1 antibody was stored at – 80 °C.

### Immunocytochemistry

Cells were permeabilized and blocked in 5% BSA, 0.3% Triton X-100 in PBS for one hour at room temperature. Antibodies were applied overnight at 4 °C (Olig2 (EMD Millipore, 1:200), NeuN (EMD Millipore, 1:200), SATB2 (Proteintech, 1:200), RARS1 (Proteintech, 1:200), S6 (Proteintech, 1:200)). The next day, cells were washed three times in PBS and secondary antibody was applied (donkey anti-rabbit, Novus, 1:500, donkey anti-mouse, Novus, 1:500). Cells were washed three times in PBS then counterstained with Hoechst 33258 for 5 min, washed twice in PBS, and mounted in Permafluor mounting medium (Invitrogen). Antibodies are listed in **Supplementary Table 9.**

### Embryonic Cortical Precursor Cultures

Cortical Precursor cultures were obtained as described by Jeong et al., 2020^49^. E14.5 embryos were obtained from the crossing of two mice heterozygous for the systemic deletion of full-length RARS1. Embryo cortices were mechanically dissociated in neurobasal medium supplemented with B27, L-glutamine, FGF, penicillin/streptomycin. The cells were seeded on PDL/laminin-coated (Sigma) coverslips at a density of 1.3 x 10^5^ cells/cm^2^ and incubated at 37 °C for 7 days. Cells were washed three times in PBS, fixed in 4% PFA for 15 mins at room temperature and stored at 4 °C in PBS until further processing.

### RNAseq library preparation and analysis

Total RNA was extracted from weighed forebrain tissue of wt/wt and dLZ/dLZ mice using the miRNeasy Mini Kit (Qiagen) according to the manufacturer’s instructions. RNA integrity and concentration were assessed prior to library preparation. Polyadenylated RNA was enriched from total RNA using the NEBNext® Poly(A) mRNA Magnetic Isolation Module (New England Biolabs). RNAseq libraries were constructed using the NEBNext® Ultra™ II RNA Library Prep with Sample Purification Beads (New England Biolabs). During adaptor ligation, the USER enzyme (New England Biolabs) was applied to resolve the hairpin-loop adaptor design, ensuring efficient downstream amplification. Finalized libraries were pooled and sequenced at the Donnelly Sequencing Centre (University of Toronto, Canada) on an Illumina NovaSeq 6000 SP platform (200-cycle flow cell), producing over 30 million paired-end reads of 100 nucleotides per sample. Downstream data processing and analysis were performed using high-performance computing infrastructure provided by Compute Ontario, SciNet, and the Digital Research Alliance of Canada.

Reads were quality controlled and adapter trimmed using Trimmomatic 0.6.4^82^ and mapped to Mus_musculus.GRCm39.114^83^ using STAR aligner 2.7.11b^84^. Counts were obtained using RSubread’s FeatureCount function (Version 2.12.3)^85^ and differentially expressed genes were identified with DESeq2 1.38.3^86^. Splicing changes were interrogated with Majiq and Voila V3^87,88^. Enrichment analysis was performed using David and Panther GO^89–91^.

### Confocal Microscopy

Confocal images were acquired on a Leica TCS SP8 microscope (Leica) using a 63×/1.40 Oil HC PL APO CS2 objective with LAS X software. Coverslips were mounted with Type F Immersion Liquid (Leica). Images were collected at 1024 × 1024 resolution with identical acquisition settings across samples, processed in ImageJ/Fiji, and exported as TIFF files.

Colocalization analysis was performed using Imaris (Oxford Instruments) on individual cell z-stacks, and Manders coefficients were calculated for each cell. Cell fluorescence was quantified using ImageJ/Fiji. Regions of interest were manually drawn around individual cells, and the integrated density was measured. Background fluorescence was determined from a cell-free area and subtracted from each measurement. Corrected total cell fluorescence (CTCF) was calculated as: CTCF = Integrated Density – (Area of selected cell X Mean fluorescence of background readings). Values were obtained for individual cells and used for statistical analysis.

### TeACAT Reagents

N-chlorosuccinimide (NCS) was purchased from Sigma. DBCO-PEG_4_-OH was purchased from ClickChem Tools (now Vector Labs). BCN-PEG_3_-TAMRA and BCN-PEG_3_-biotin sulfone were made in-house using published procedures^56^. All PBS used is Dulbecco’s phosphate buffered saline without calcium or magnesium. Streptavidin-agarose used for proteomics was purchased from Genscript. MS-grade trypsin was purchased from Promega.

### Administration of TePhe and Tissue Collection

TePhe was synthesized using a reported method^92^. TePhe was formulated at 12 mg/mL in 40% Captisol in PBS, sterile-filtered through a 0.22 µm PES syringe filter, and stored at -80 °C until use. Mice were dosed twice with 45 mg/kg TePhe (between 75 and 100 µL per IP injection) 4 hours apart; vehicle controls received equivalent volumes of 40% Captisol in PBS. Animals were euthanized by CO_2_ inhalation 8 hours after the initial injection. Olfactory bulb and forebrain tissues were dissected separately, immediately frozen in liquid nitrogen, and stored at -80 °C. The remainder of the brain was fixed by immersion in 4% paraformaldehyde in PBS for 3 days, followed by routine procedures for dehydration, paraffin infiltration and embedding. Formalin-fixed, paraffin-embedded (FFPE) brains were coronally sectioned to 4 µm using a Leica microtome (Centre for Immune Analytics, University of Toronto) and affixed to SuperFrost glass slides.

### TePhe detection in FFPE brain sections

4 µm-thick coronal sections on glass slides were baked at 60 °C for 30 min, followed by dewaxing in xylenes (2 x 5 min) and rehydration in graded alcohols (100%, 90%, 80%, 70%, each 2 x 5 min) and deionized water (5 min). Slides were equilibrated in alkylation buffer (100 mM Tris + 100 mM NaCl, pH 8.5 + 0.025% Triton X-100) for 5 min, followed by capping of free thiols in fresh alkylation buffer containing 10 mM iodoacetamide in the dark for 20 min. Slides were rinsed in PBS, then equilibrated in OSTAC reaction buffer (10 mM NaOAc, pH 5 + 0.025% Triton X-100) for 5 min. Slides were transferred to OSTAC blocking solution (150 µM DBCO + 500 µM NCS in OSTAC reaction buffer, prepared immediately before use) and incubated for exactly 10 min. Slides were then transferred to OSTAC labeling solution (1 µM BCN-PEG_4_-TAMRA + 100 µM NCS in OSTAC reaction buffer, prepared immediately before use) and incubated for exactly 10 min. To stop the reaction, slides were quickly transferred into an excess volume of washing buffer (PBS + 0.025% Triton X-100). This was followed by immersion in fresh washing buffer and two further washes in PBS. Finally, samples were stained with DAPI by immersion in 2.5 µg/mL solution in PBS for 20 min, followed by two brief PBS washes and mounting in Prolong Gold antifade mounting medium. Images were obtained on a Leica Mica epifluorescence microscope with spectral deconvolution (10x objective dry objective). Images were processed in ImageJ, with adjustment of raw DAPI and TAMRA channel intensities by a common factor (multiplication 20-fold, followed by subtraction of 3000 AU) to obtain optimal intensities for viewing.

### Global protein synthesis measurements by TePhe-based fluorescence labeling

Unless otherwise stated, steps were performed at room temperature. Frozen olfactory bulb tissues were briefly thawed at ambient temperature, roughly crushed using a pipet tip, suspended in lysis buffer (200-500 µL of PBS + 0.5% SDS depending on amount of tissue), and homogenized by repeatedly pulling the suspension through a 22-gauge hypodermic needle. Lysates were quantified by BCA assay (Pierce). 30 µL lysates (diluted in PBS + 0.5% SDS to 1.1 mg/mL protein) were used for the labeling reaction. EDTA was added (final conc. 3 mM) and the mixture incubated for 15 min. Free thiols were alkylated by direct addition of iodoacetamide (10 mM) and incubation for 20 min in the dark, after which samples were directly subjected to OSTAC chemistry. The OSTAC labeling sequence consisted of addition of DBCO-PEG_4_-OH (150 µM) and NCS (600 µM) and incubation for exactly 10 min, followed by further addition of BCN-PEG_3_-TAMRA (10 µM) and NCS (100 µM) and incubation for 10 min. Reactions were quenched by addition of 5x gel loading dye (300 mM Tris-Cl pH 6.8, 50 w/v % glycerol, 10% w/v SDS, 20% v/v β-mercaptoethanol, trace bromophenol blue). Samples were boiled for 2 min, the entire volume loaded onto a 1.5 mm thickness 10% SDS-PAGE gel, and fractionated at 125-130 V for 1 hr. TAMRA fluorescence was measured under green LED illumination on a Syngene G:Box Chemi XT 4 gel documentation system with the gel still within its glass backing. Routine Coomassie staining (1.5 hr), destaining (overnight) and imaging were then performed. Densitometry analysis was performed in GelAnalyzer to normalize TAMRA of each lane to Coomassie intensity.

### TeACAT Proteomics

Frozen forebrain tissues were briefly thawed at ambient temperature, roughly crushed using a pipet tip, suspended in lysis buffer (500 µL PBS + 1% SDS per sample), and homogenized by pulling the suspension through a 22-gauge hypodermic needle, then passed through a 70 µm mesh cell strainer. Protein concentrations were determined by BCA assay and all samples diluted in PBS + 1% SDS to match the lowest concentration sample. 400 µg input per sample were further diluted by addition of alkylation buffer (220 mM Tris pH 5, 220 mM NaCl) to a final concentration of ∼0.5 mg/mL protein in buffer containing 100 mM Tris, 100 mM NaCl and 0.5% SDS. Reduction and alkylation were performed by addition of DTT (final conc. 7 mM) and incubation at 55 °C for 45 min, cooling for 5 min, addition of chloroacetamide (14 mM) and incubation at room temperature for 25 min in the dark, and quenching with additional DTT (7 mM). Reaction mixtures were divided into 2 x 2 mL microcentrifuge tubes and precipitated by addition of 4 volumes of acetone and left at -20 °C overnight. Precipitates were pelleted at 18000 x g for 5 min, supernatant was discarded, and the pellet resuspended in fresh acetone (750 µL per microcentrifuge tube) by brief sonication in a water bath. Acetone suspension aliquots of the same sample were recombined for subsequent washes. Centrifugation, supernatant removal, and resuspension steps were repeated for a total of 4 acetone washes to fully remove DTT and chloroacetamide derivatives. Pellets were briefly air-dried and redissolved in PBS + 0.5% SDS (300 µL) with end-to-end rocking for 2 hours. 320 µg input at 1.0 mg/mL protein in PBS + 0.5% SDS was subjected to OSTAC labeling. DBCO-PEG_4_-OH (150 µM) + NCS (500 µM) were added and the mixture incubated for exactly 10 min, followed by BCN-PEG_3_-Biotin sulfone (10 µM) + NCS (100 µM) and incubation for exactly 10 min. The reaction was quenched by addition of DTT (5 mM), and precipitated by addition of 4 volumes of acetone and left overnight at -20 °C. Precipitates were collected by centrifugation at 18000 x g for 5 min and the pellet resuspended in fresh acetone (1.5 mL) by brief sonication in a water bath. This process was repeated for a total of 4 acetone washes to remove excess probe and small molecules from the OSTAC step. Pellets were briefly air-dried, redissolved in PBS + 1% SDS (700 µL) and quantified by BCA assay. Samples were diluted to 0.2 mg/mL in PBS + 1% SDS for streptavidin capture.

Streptavidin-agarose beads were equilibrated by washing twice in PBS and 4 times in PBS + 1% SDS (180 µL slurry, wash volume = 1.5 mL). Each wash consisted of 3 cycles of aggressive vortexing of the bead-buffer mixture for 5 sec followed by inversion of the tube 5 times. After the final wash, beads were aliquoted (16 µL slurry or ∼8 µL settled beads per tube). Samples were added (290 µg input per sample, with the exception of one wt sample at 250 µg) and the mixtures placed on over-the-end rotation for 2 hours at room temp. Bead washing consisted of centrifugation at 2000 x g for 1 min, removal of supernatant, addition of fresh PBS + 1% SDS (1.4 mL), and mixing (4 cycles of vortexing for 5 sec and inversion of the tube 5 times, then placement on over-the-end rotator for 5 min). Washing was repeated for a total of 4 times in PBS + 1% SDS, 2 times in PBS only, and 2 times in 100 mM ammonium bicarbonate, pH 8. Centrifugation times were increased following later washes as differential swelling in various buffers led to decreased ease of pelleting streptavidin-agarose. The supernatant was removed and beads suspended in 95 µL fresh 100 mM ammonium bicarbonate. Sequencing grade trypsin (Promega) was added at 2.4 µg per sample and the mixture digested on a thermomixer at 37 °C, 1250 rpm overnight. After 16 h, beads were pelleted and 100 µL of supernatant were transferred to a new tube. Digestion was quenched by addition of 1% formic acid followed by 1 h incubation at room temp. Digests were stored at -80 °C until C18 clean-up. Clean-up was performed using C18 stage tips (Pierce #87784) according to manufacturer’s instructions, but with formic acid in place of trifluoroacetic acid. Eluted peptides were dried under vacuum and reconstituted in MS-grade water + 0.1% formic acid (8 µL). 1 µL sample was taken for BCA quantitation. Samples were diluted as needed to achieve the target injection concentration of 800 ng peptides per 5.5 µL.

### Co-immunoprecipitation

Brain tissue was homogenized in TBS with% NP-40 and 1 mM MgCl_2_ supplemented with protease and phosphatase inhibitors. The homogenized mixture was incubated with gentle agitation for 30 min at 4°C followed by centrifugation at 14,000 x*g* for 20 min to remove insoluble components. Lysates were quantified by BCA assay and equal amounts of total protein were added to 30 μl of pre-equilibrated protein G beads (Genscript) along 2 μg of anti-RARS1 antibody (homemade, verified in ^55^, sequence published in ^81^). After 12 h, beads were washed three times with 500 μl of TBS. Beads were then stored in PBS (Gibco) at −80 °C. Peptides were eluted from protein G beads by trypsin digest at room temperature overnight (16 h) while shaking (5 ng/μl trypsin, 2 M urea, 50 mM Tris–HCl, pH 7.5, 0.2 mM DTT, and 1 mM chloroacetamide). Digested peptides were desalted with C18 tips (Pierce) according to the manufacturer’s instructions. Peptides were eluted in a final volume of 30 μl, dried using a SpeedVac (Thermo Fisher), and resuspended in 8 μL of 0.1% formic acid in MS-grade water.

### LC-MS/MS acquisition

For both TeACAT and co-immunoprecipitation experiments, eluted peptides were dried under vacuum and reconstituted in MS-grade water + 0.1% formic acid (8 µL). 1 µL sample was taken for BCA quantitation. Samples were diluted as needed to achieve the target injection concentration of 800 ng peptides per 5.5 µL. Peptides were separated using a 14 cm PicoFrit analytical column (75 μm inner diameter) packed in-house with ReproSil-Pur C18-AQ resin (1.9 μm particle size) and operated at a flow rate of 300 nL/min. Liquid chromatography was performed on an EASY-nLC 1200 system (Thermo Fisher Scientific) coupled to a Q Exactive HF-X mass spectrometer (Thermo Fisher Scientific). Mobile phase A consisted of 0.1% formic acid, while mobile phase B consisted of 80% acetonitrile containing 0.1% formic acid. Peptides were eluted using a linear gradient of 2-5% B over 1 min, 5-26% B over 70 min, 26-60% B over 19 min, and 60-100% B over 15 min, followed by a 15 min wash at 100% B.

Mass spectrometric data were acquired in positive ion mode using a data-dependent acquisition method, in which the 10 most intense precursor ions from each full scan were selected for fragmentation. Full MS scans were acquired over an *m/z* range of 300-1650 at a resolution of 60,000, with an automatic gain control (AGC) target of 1 × 10^6^ ions. MS/MS spectra were generated by higher-energy collisional dissociation (HCD) using a normalized collision energy of 27 and acquired at a resolution of 15,000 (*m/z* 200). Precursor ions were isolated with a 1.4 *m/z* isolation window and accumulated for a maximum injection time of 25 ms or until an AGC target of 1 × 10^5^ ions were reached. Singly charged ions and ions with unassigned charge states were excluded from fragmentation. Dynamic exclusion was enabled with a duration of 15 s to prevent repeated sequencing of the same precursor ions.

### Proteomics Data Analysis

For TeACAT: Searches were performed in MaxQuant^93^ against the mouse reference proteome (UP000000589) with carbamidomethyl-Cys set as a fixed modification and N-terminal acetylation, Met oxidation (thioether to sulfoxide), and Trp oxidation (indole to oxindole) set as variable modifications. Protein groups registering a minimum ratio count of 2 were included for label free quantification. Data processing was performed using Perseus^94^ software. Enrichment analysis was performed using ShinyGO^95^.

For co-immunoprecipitation: MS data were processed with MaxQuant 2.6.4.0 and Perseus 2.1.2^94^ (both were developed at the Max Planck Institute of Biochemistry) to identify RARS1 interaction partners as described in ^21^. In brief, raw files were searched against the *Mus musculus* reference proteome UP000000589 (UniProt^96^) with the Andromeda search engine integrated into MaxQuant: M oxidation, N-terminal acetylation as variable; cysteine carbamidomethyl as fixed modification; and two missed cleavages maximum. First search peptide tolerance = 20 ppm; main search peptide tolerance = 4.5 ppm; protein false discovery rate = 0.1; and minimum peptides = 1. Default settings were used for label-free quantification (minimum ratio count = 2). The resulting protein groups file was loaded into Perseus and filtered for “reverse”, “potential contaminants,” and “only identified by site.” The log_2_ values of label-free quantification intensities were calculated, samples were grouped according to experimental conditions (wt/wt, dLZ/dLZ), and all proteins with less than two valid values/group were discarded. Missing values were replaced from a normal distribution (width 0.3 and downshift 1.8), and a two-tailed unpaired Student’s *t* test was used to calculate *t* test significance and difference (fold change) for each protein by comparing wt versus RARS1 dLZ interactomes.

### Statistics

Significantly enriched interactors and differentially synthesized proteins were identified using unpaired, two-tailed Students t-test in Perseus. Differentially expressed genes were determined using DESeq2, padj < 0.05. Differentially spliced genes were identified using Majiq/Voila, with sites filtered for a change in percent spliced in (PSI) >5% and a *p*-value <0.05. Unpaired Student’s t-test or One-way ANOVA calculated with GraphPad Prism (GraphPad Software Inc.) were used for all other data. Replicates are individual animals unless stated otherwise. Littermates were used as controls whenever possible.

## Acknowledgements

We are deeply grateful to the Bioscience Support Facility for mouse housing, husbandry, support, and training. We thank the Cell and Systems Biology Imaging core for access to confocal microscopy and Imaris as well as the Centre for Applied Genomics and the Donnelly Sequencing core for next generation sequencing. We also thank the Transgenic Core facility at the Salk Institute for Biological Studies and Dr. Paul Schimmel for support during the generation of the initial Rars1^loxP/loxP^ mouse line. We thank Felicia Pais Araújo, Chloe Taylor, and Eaarad Aftab for assistance with mouse genotyping as well as Emad Naimi Ghahroodi for assistance with counting primary oligodendrocytes.

## Author contribution

Conceptualization: SPN, HC. Investigation: SPN, RT, AH, YJB, QS, SV, DH, FLA, CMM, JZ, HG, JC, SW, ND. Methodology: SPN, RT, AH, YJB, QS, DH, SW. Resources: AW, MN, VM, SAY, HC. Supervision: AW, MN, VEM, SAY, HC. Visualization: SPN, RT, AH, YJB, QS, HC. Writing – Original Draft Preparation: SPN, HC. Writing – Review & Editing: all authors.

