## Extended Data Figures for "Aminoacyl-tRNA synthetase organization is crucial for mammalian brain development"

### Supplementary Data:

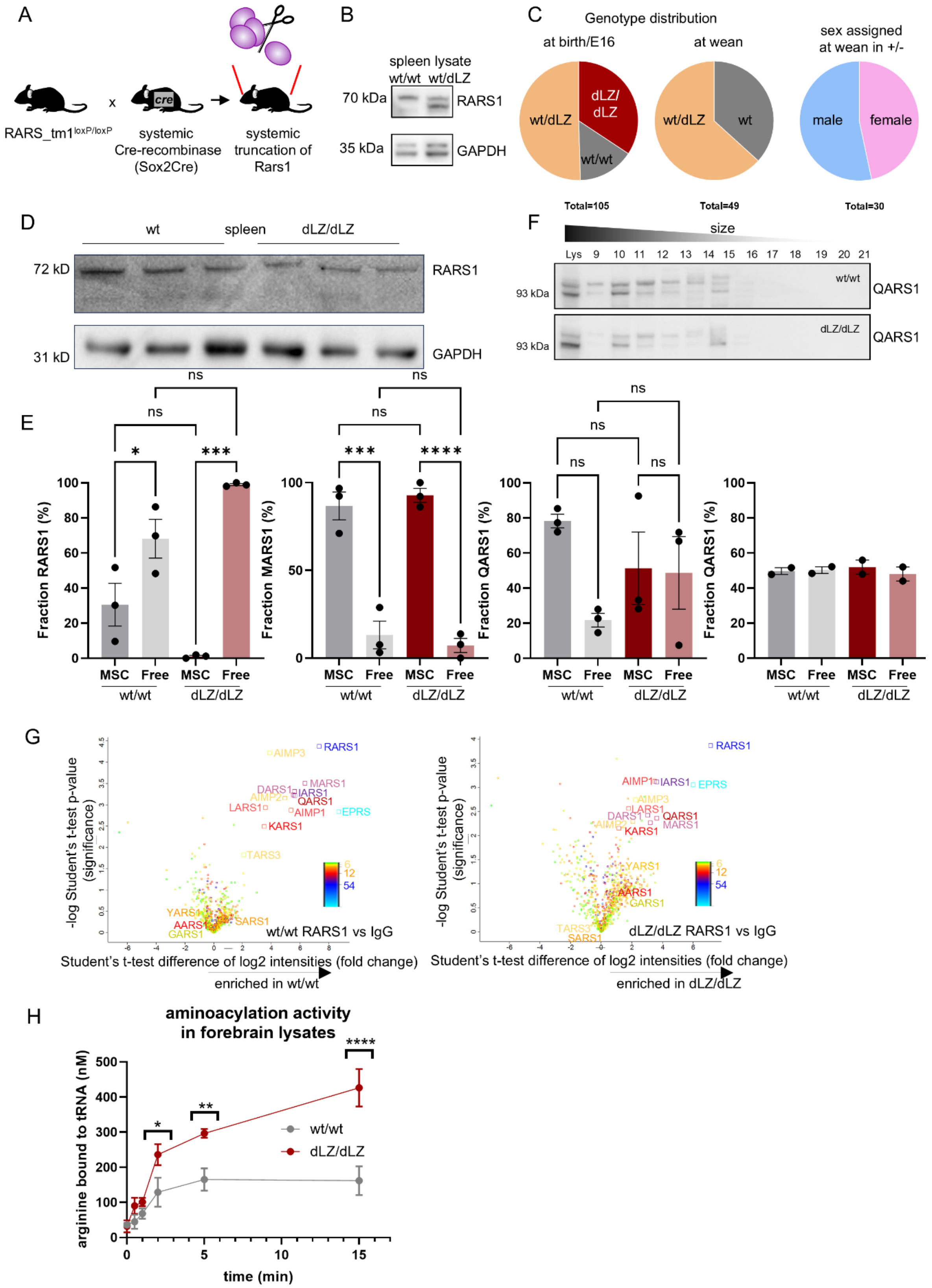
A
RARS_tm1loxP/loxP
x
systemic Cre-recombinase (Sox2Cre)
systemic truncation of Rars1
B
spleen lysate
wt/wt wt/dLZ
70 kDa
RARS1
35 kDa
GAPDH
C
Genotype distribution
at birth/E16
at wean
sex assigned at wean in +/-
wt/dLZ
dLZ/dLZ
wt/wt
wt
male
female
Total=105
Total=49
Total=30
D
wt
spleen
dLZ/dLZ
72 kD
RARS1
31 kD
GAPDH
F
size
Lys 9 10 11 12 13 14 15 16 17 18 19 20 21
93 kDa
wt/wt
QARS1
dLZ/dLZ
E
Fraction RARS1 (%)
Fraction MARS1 (%)
Fraction QARS1 (%)
MSC Free MSC Free
wt/wt dLZ/dLZ
ns
*
***
****
G
-log Student's t-test p-value (significance)
Student's t-test difference of log2 intensities (fold change)
wt/wt RARS1 vs IgG
enriched in wt/wt
dLZ/dLZ RARS1 vs IgG
enriched in dLZ/dLZ
RARS1 AIMP3 MARS1 DARS1 IARS1 AIMP2 QARS1 LARS1 AIMP1 EPRS KARS1 TARS3 YARS1 AARS1 SARS1 GARS1
H
aminoacylation activity in forebrain lysates
arginine bound to tRNA (nM)
time (min)
wt/wt
dLZ/dLZ

**Extended Data Fig. 1: Systemic deletion of RARS1 LZ and assessment of MSC integrity**

A) Overview of breeding scheme leading to systemic RARS1 N-terminal truncation. Sox2 is active in epiblasts, therefore all epiblast-derived cells lack exon 2 in RARS1. Following F1, mice were bred in wt/dLZ x wt/dLZ matings. B) Immunoblot of spleen lysate showing the increased expression of N-terminally truncated RARS1 in heterozygous mice. GAPDH was used as a loading control. C) Genotype and sex distribution. Left: Genotype was assigned either at birth or in E16 embryos. dLZ/dLZ embryos and pups were found at slightly above Mendelian ratios. Middle: dLZ/dLZ mice were absent at wean. Close observations of newborn pups suggested that they succumbed shortly after birth. Right: Sex distribution in wt/dLZ heterozygous animals.

D) Immunoblot of RARS1 wt/wt and dLZ/dLZ (*Emx1*-Cre+ x Rars1$^{loxP/loxP}$) spleen lysates showing only full-length RARS1 in spleen, confirming the tissue-specificity of exon 2 deletion. E) Alternative QARS1 blot with a second antibody following size exclusion chromatography. F) Densitometric quantification of aminoacyl-tRNA synthetase from size exclusion chromatography brain lysate fractionation (see Figure 1D for blots). SEC fractions were individually quantified, and values were added to 100%. Fractions attributed to the multisynthetase complex (fractions 9-11) and fractions corresponding to free-standing aminoacyl-tRNA synthetases (all later fractions) are shown as percentages. RARS1: n=3, 3, 3, 3. Ordinary, one-way ANOVA, multiple comparisons, *p = 0.0476, p > 0.05, ***p = 0.0001, p > 0.05. MARS1: n=3, 3, 3, 3. Ordinary, one-way ANOVA, multiple comparisons, ***p = 0.002, ns p > 0.05, ****p < 0.0001, ns p > 0.05. QARS1 (Proteintech) n=3, 3, 3, 3. Ordinary, one-way ANOVA, multiple comparisons, p > 0.05, p > 0.05, p > 0.05, p> 0.05. QARS1: (ThermoFischer) n=2, 2, 2, 2. G) Left panel: RARS1 wt/wt interactome (anti-RARS1 antibody pulldown compared with IgG control pulldown in wt/wt brain lysates). Right panel: RARS1 dLZ/dLZ interactome (anti-RARS1 antibody pulldown compared with IgG control pulldown in dLZ/dLZ brain lysates). Color coding denotes the number of peptides. n = 3, 3. H) Aminoacylation activity kinetics in forebrain lysates of *Emx1*-Cre+ x Rars1$^{loxP/loxP}$ animals. n=3, 3. Unpaired, two-tailed t test, *p=0.0224, **p=0.0027, **p=0.0024.

B, C). wt: wildtype. dLZ: RARS1 delta leucine zipper, mice originated from *Sox2* x Rars1$^{loxP/loxP}$ and were bred as wt/dLZ animals following F1.

D-H) wt/wt: wildtype RARS1, *Emx1*-Cre+ x Rars1$^{wt/wt}$. dLZ/dLZ: RARS1 delta leucine zipper, *Emx1*-Cre+ x Rars1$^{loxP/loxP}$. Replicates are individual animals. All experiments were performed in 2 months old male mice.

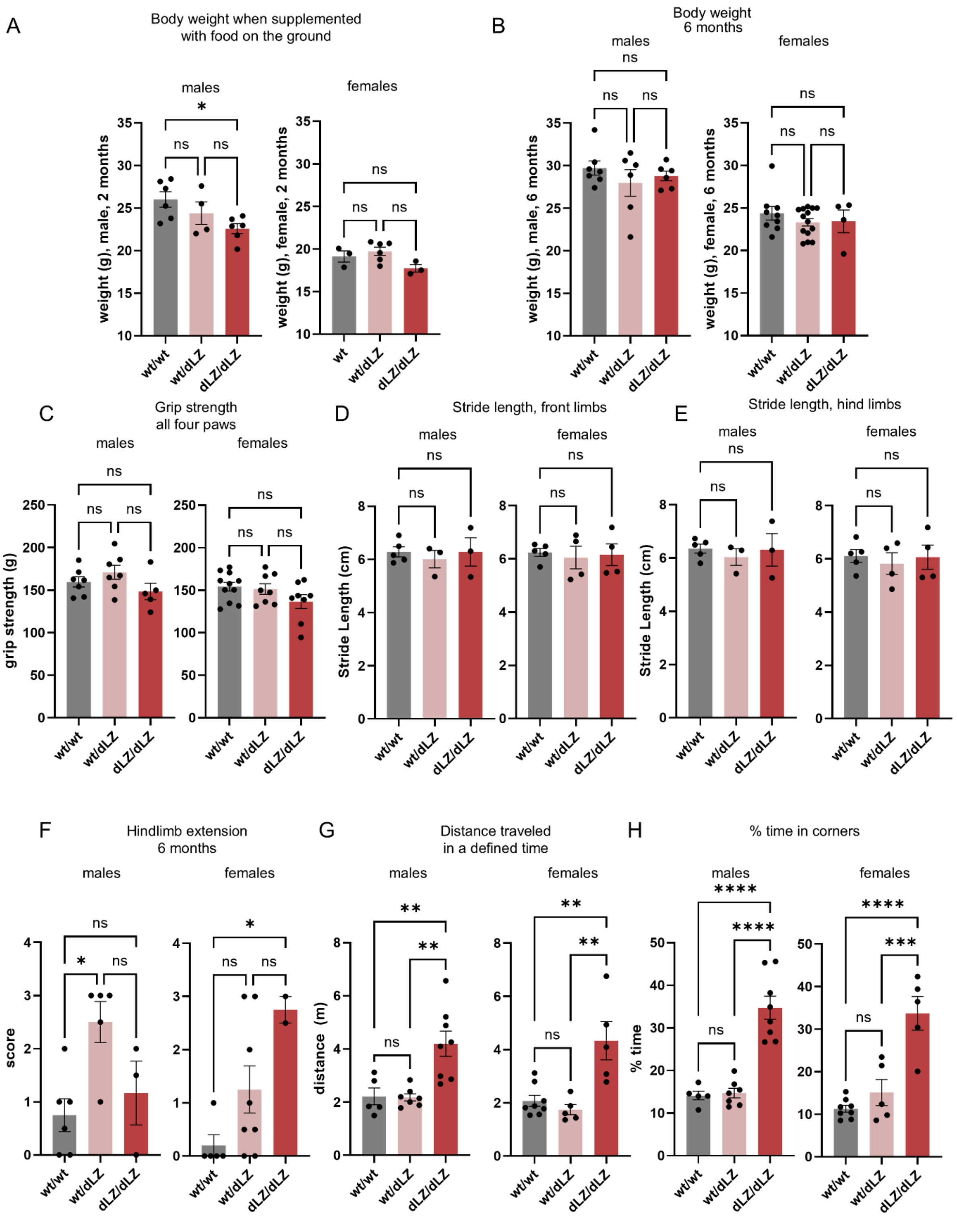
A
Body weight when supplemented with food on the ground
males
females
weight (g), male, 2 months
weight (g), female, 2 months
B
Body weight 6 months
males
females
weight (g), male, 6 months
weight (g), female, 6 months
C
Grip strength all four paws
males
females
grip strength (g)
D
Stride length, front limbs
males
females
Stride Length (cm)
E
Stride length, hind limbs
males
females
Stride Length (cm)
F
Hindlimb extension 6 months
males
females
score
G
Distance traveled in a defined time
males
females
distance (m)
H
% time in corners
males
females
% time
wt/wt
wt
wt/dLZ
dLZ/dLZ
ns
*
**
***
****

**Extended Data Fig. 2: Additional phenotype characterization of mice with N-terminally truncated RARS1.**

A) Body weight of ground fed mice. Males: n = 8, 5, 8. Ordinary, one-way ANOVA, multiple comparisons, ns p > 0.05, **p = 0.0047. Females: n = 3, 6, 3. Ordinary, one-way ANOVA, multiple comparisons, ns p > 0.05. B) Body weight of mice at 6 months of age. Males: n = 7, 6, 6. Females: n = 9, 14, 4. Ordinary, one-way ANOVA, multiple comparisons, ns p > 0.05. C) Grip strength measured by attachment of the mouse to a metal grid connected to a grip strength meter. The mouse was allowed to attach itself with all four paws. Males: n=7, 7, 5. Females: n=11, 8, 8. Ordinary, one-way ANOVA, multiple comparisons, ns p > 0.05. D) Stride length of the front limbs, measured by applying ink to the front paws and measuring footprints on grid paper. Males: n=5, 3, 3. Females: n=5, 4, 4. Ordinary, one-way ANOVA, multiple comparisons, ns p > 0.05. E) Stride length of the hind limbs, measured by applying ink to the back paws and measuring footprints on grid paper. Males: n=5, 3, 3. Females: n=5, 4, 4. Ordinary, one-way ANOVA, multiple comparisons, ns p > 0.05. F) Hindlimb extension test to score abnormalities in motor neuron function and limb clasping. The higher the score, the worse the animal performed. Males: n=6, 5, 3. Females: n=5, 8, 2. Ordinary, one-way ANOVA, multiple comparisons, *p = 0.0156, *p = 0.0283, ns p > 0.05. G) Total distance traveled was measured through deep learning-based video analysis. Males: n=5, 7, 8. Ordinary, one-way ANOVA, multiple comparisons, *p = 0.0054, *p = 0.0021. Females: n=8, 5, 5. Ordinary, one-way ANOVA, multiple comparisons, **p = 0.0021, **p = 0.0016. H) Percentage of time spent in the corner of the open field transparent box. Males: n=5, 7, 8. Ordinary, one-way ANOVA, multiple comparisons, ****p < 0.0001. Females: n=8, 5, 5. Ordinary, one-way ANOVA, multiple comparisons, ****p < 0.0001.

A-H) wt/wt: wildtype RARS1, *Emx1*-Cre+ x Rars1$^{wt/wt}$. dLZ/dLZ: RARS1 delta leucine zipper, *Emx1*-Cre+ x Rars1$^{loxP/loxP}$. Replicates are individual animals.

A, C-E, G, H) All experiments were performed in 2 months old male mice.

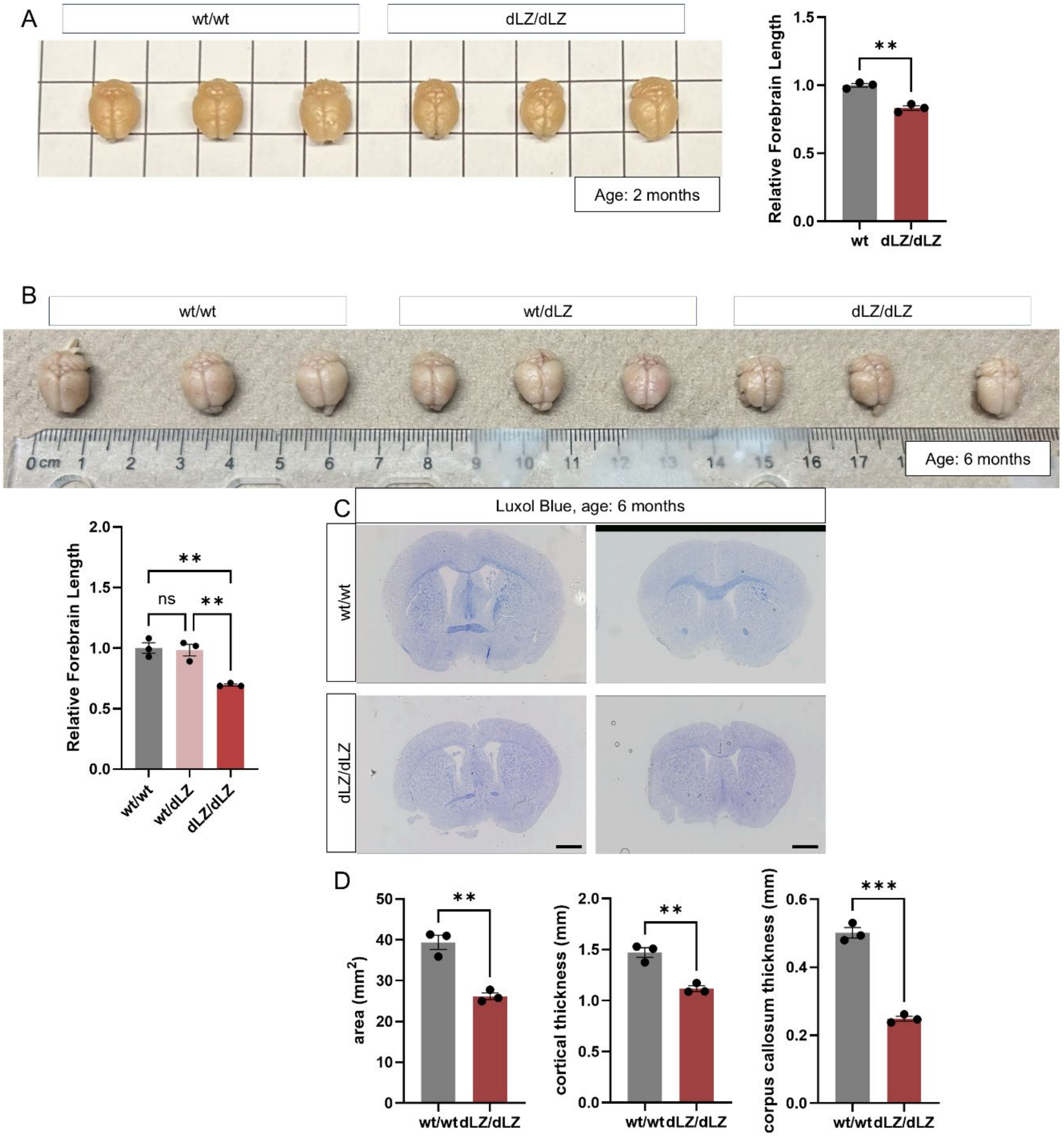
A
wt/wt
dLZ/dLZ
Age: 2 months
Relative Forebrain Length
1.5
1.0
0.5
0.0
**
wt
dLZ/dLZ
B
wt/wt
wt/dLZ
dLZ/dLZ
Age: 6 months
0 cm 1 2 3 4 5 6 7 8 9 10 11 12 13 14 15 16 17
Relative Forebrain Length
2.0
1.5
1.0
0.5
0.0
**
ns
**
wt/wt
wt/dLZ
dLZ/dLZ
C
Luxol Blue, age: 6 months
wt/wt
dLZ/dLZ
D
area (mm2)
50
40
30
20
10
0
**
wt/wt
dLZ/dLZ
cortical thickness (mm)
2.0
1.5
1.0
0.5
0.0
**
wt/wt
dLZ/dLZ
corpus callosum thickness (mm)
0.6
0.4
0.2
0.0
***
wt/wt
dLZ/dLZ

**Extended Data Fig. 3: Reduced forebrain volume in mice with N-terminally truncated RARS1 at 6 months of age.**

A) Picture of whole mount brains from 2-month-old female mice. Grid size is 1 cm. n=3, 3. Unpaired, two-tailed t test, **p=0.0014. B) Picture of whole mount brains taken from 6-month-old male mice. Quantification of relative forebrain size. n=3, 3, 3. Ordinary, one-way ANOVA, multiple comparisons, **p = 0.0030, **p = 0.0040. C) Cryosections of mouse brains collected at 6 months, stained with Luxol fast blue, which visualizes myelin. Bar=1 mm. D) Total area, cortical thickness, and corpus callosum thickness quantified from matched serial sections of mouse forebrains obtained from 6 months old male mice. n=3, 3. Unpaired, two-tailed t test, **p = 0.0024, **p = 0.0031, ***p = 0.0001.

A-D) wt/wt: wildtype RARS1, *Emx1*-Cre+ x Rars1$^{wt/wt}$. dLZ/dLZ: RARS1 delta leucine zipper, *Emx1*-Cre+ x Rars1$^{loxP/loxP}$. Replicates are individual animals.

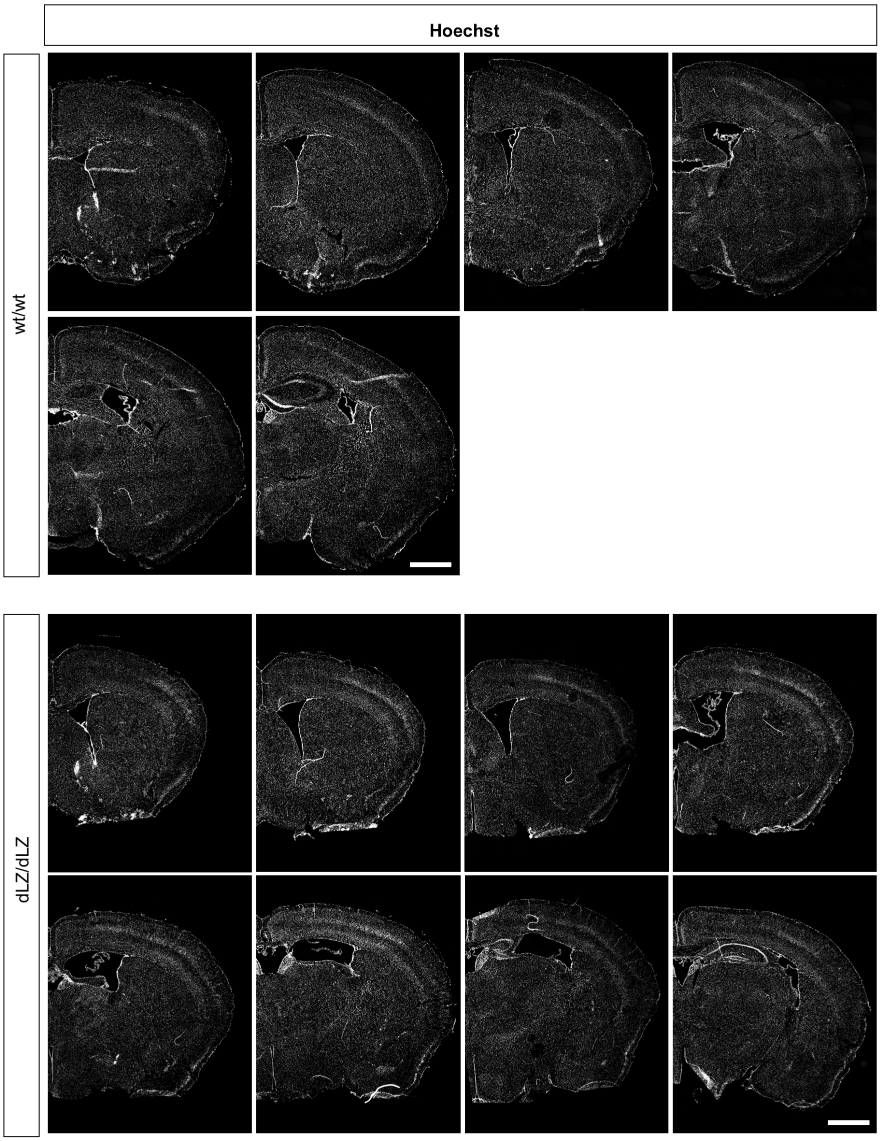
Hoechst
wt/wt
dLZ/dLZ

**Extended Data Fig. 4: Nuclear Staining in wild-type and dLZ/dLZ brains.**

Nuclear staining in hemicoronal sections from wt and dLZ/dLZ animals highlights overall anatomical features. 7 to 8 levels of sections are shown. Levels 1 to 3 correspond to the morphological quantifications in Figure 3. 18 μm thick sections were obtained. Sections are approximately 360 μm apart. Scale bar = 1000 μm.

A-C) wt/wt: wildtype RARS1, *Emx1*-Cre+ x $Rars1^{wt/wt}$. dLZ/dLZ: RARS1 delta leucine zipper, *Emx1*-Cre+ x $Rars1^{loxP/loxP}$. Replicates are individual animals.

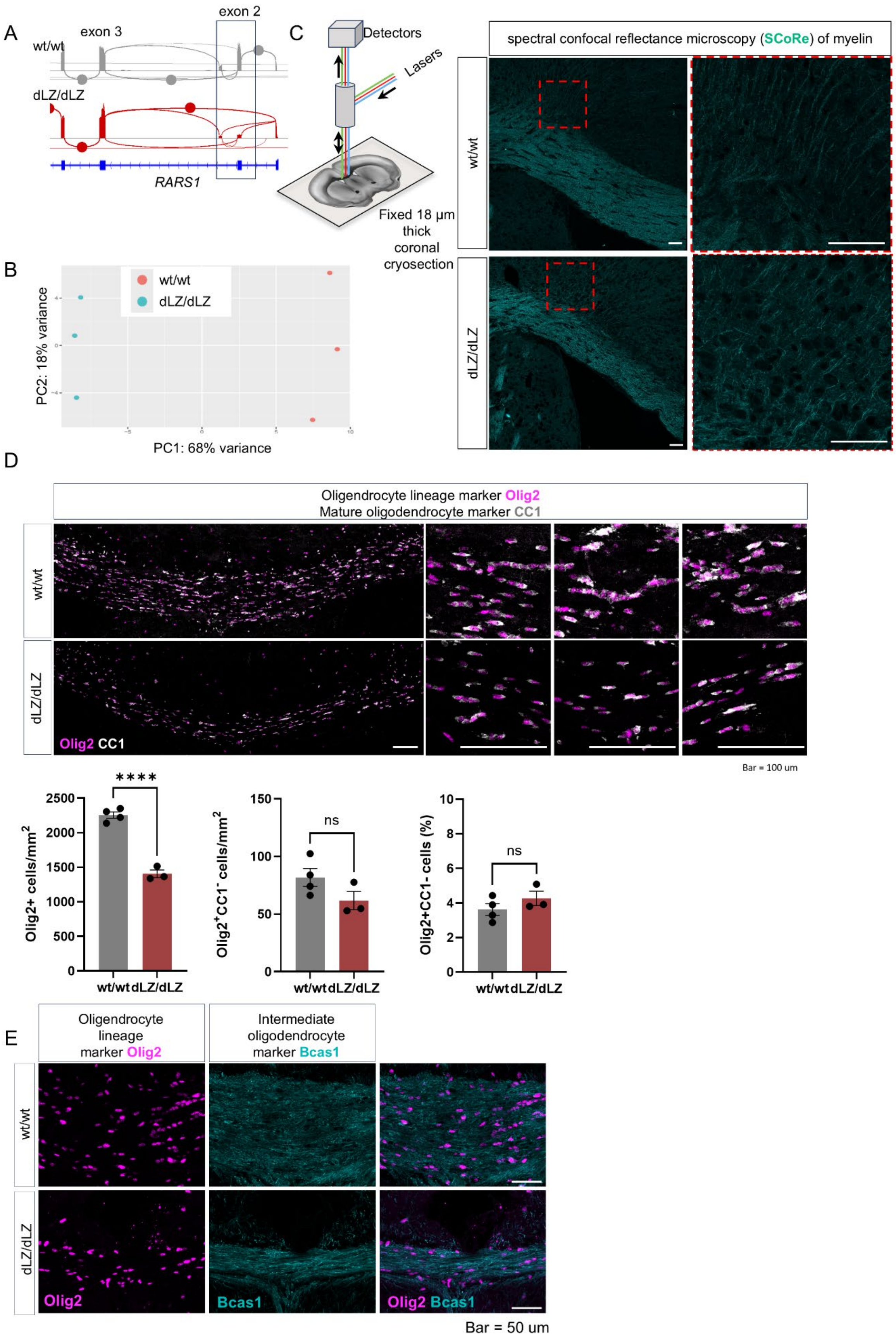

A
exon 2
exon 3
wt/wt
dLZ/dLZ
RARS1
C
Detectors
Lasers
Fixed 18 µm thick coronal cryosection
spectral confocal reflectance microscopy (SCoRe) of myelin
wt/wt
dLZ/dLZ
B
wt/wt
dLZ/dLZ
PC2: 18% variance
PC1: 68% variance
D
Oligendrocyte lineage marker Olig2
Mature oligodendrocyte marker CC1
wt/wt
dLZ/dLZ
Olig2 CC1
Bar = 100 um
****
Olig2+ cells/mm2
wt/wt dLZ/dLZ
ns
Olig2+CC1- cells/mm2
ns
Olig2+CC1- cells (%)
E
Oligendrocyte lineage marker Olig2
Intermediate oligodendrocyte marker Bcas1
Olig2
Bcas1
Olig2 Bcas1
Bar = 50 um

**Extended Data Fig. 5: RNAseq, SCoRE imaging, and additional images comparing myelin and oligodendrocyte lineage cells**

A) Sashimi plot showing altered exon usage in RARS1 dLZ/dLZ mouse forebrains due to Cre-driven recombination at loxP sites flanking RARS1 exon 2. B) Principal component analysis suggested differential expression between genes in wt and dLZ/dLZ animals. C) Simplified schematic illustrating the principle of SCoRE and representative image of SCoRe images acquired. D) Representative overview of the region of the corpus callosum quantified for $Olig2^+$ (Oligodendrocyte transcription factor 2, oligodendrocyte lineage marker) and $CC1^+$ (mature oligodendrocyte marker, Quakin 7) nuclei. Bar = 100 µm. Number of oligodendrocyte-lineage cells Olig2 /mm$^2$ in wt/wt and dLZ/dLZ mice. n = 4, 3. Unpaired, two-tailed t test, ****$p < 0.0001$. Number of immature oligodendrocyte cells (Olig2+CC1-) /mm$^2$ in wt/wt and dLZ/dLZ mice. n = 4, 3. Unpaired, two-tailed t test, $p > 0.05$. Percentage of immature oligodendrocyte-lineage cells (Olig2+, CC1- per Olig2+). n = 4, 3. Unpaired, two-tailed t-test, $p>0.05$. E) Immunofluorescence of Olig2 and newly generated myelin, indicated by Bcas1 (Breast carcinoma-amplified sequence 1 homolog, marker of intermediate oligodendrocytes) in forebrain sections. Bar = 50 µm.

A-E) wt/wt: wildtype RARS1, *Emx1*-Cre+ x Rars1$^{wt/wt}$. dLZ/dLZ: RARS1 delta leucine zipper, *Emx1*-Cre+ x Rars1$^{loxP/loxP}$. Replicates are individual animals.

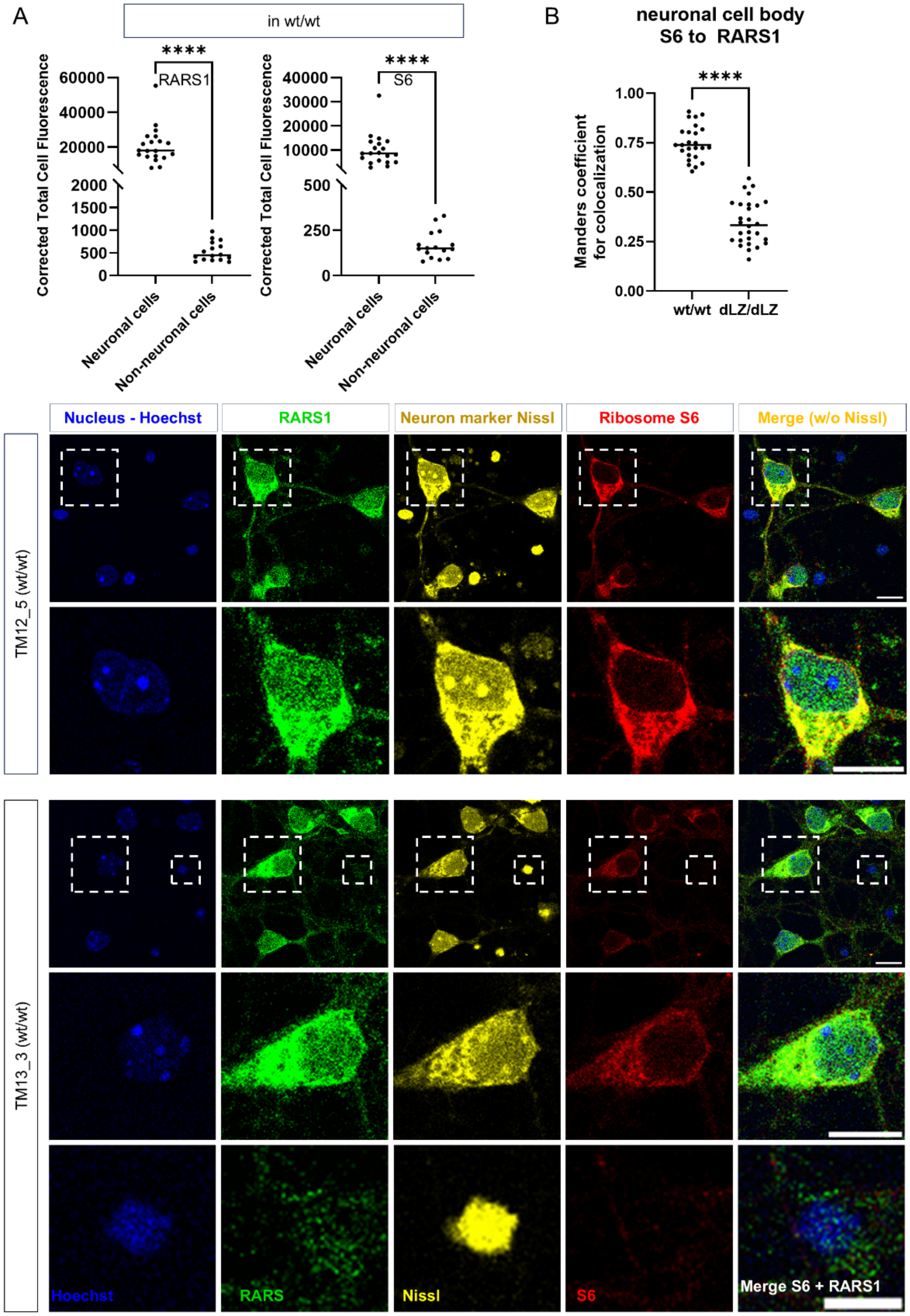

A
in wt/wt
****
RARS1
Corrected Total Cell Fluorescence
60000
40000
20000
2000
1500
1000
500
0
Neuronal cells
Non-neuronal cells
****
S6
40000
30000
20000
10000
500
250
0
B
neuronal cell body
S6 to RARS1
****
Manders coefficient for colocalization
1.00
0.75
0.50
0.25
0.00
wt/wt
dLZ/dLZ
Nucleus - Hoechst
RARS1
Neuron marker Nissl
Ribosome S6
Merge (w/o Nissl)
TM12_5 (wt/wt)
TM13_3 (wt/wt)
Hoechst
RARS
Nissl
S6
Merge S6 + RARS1

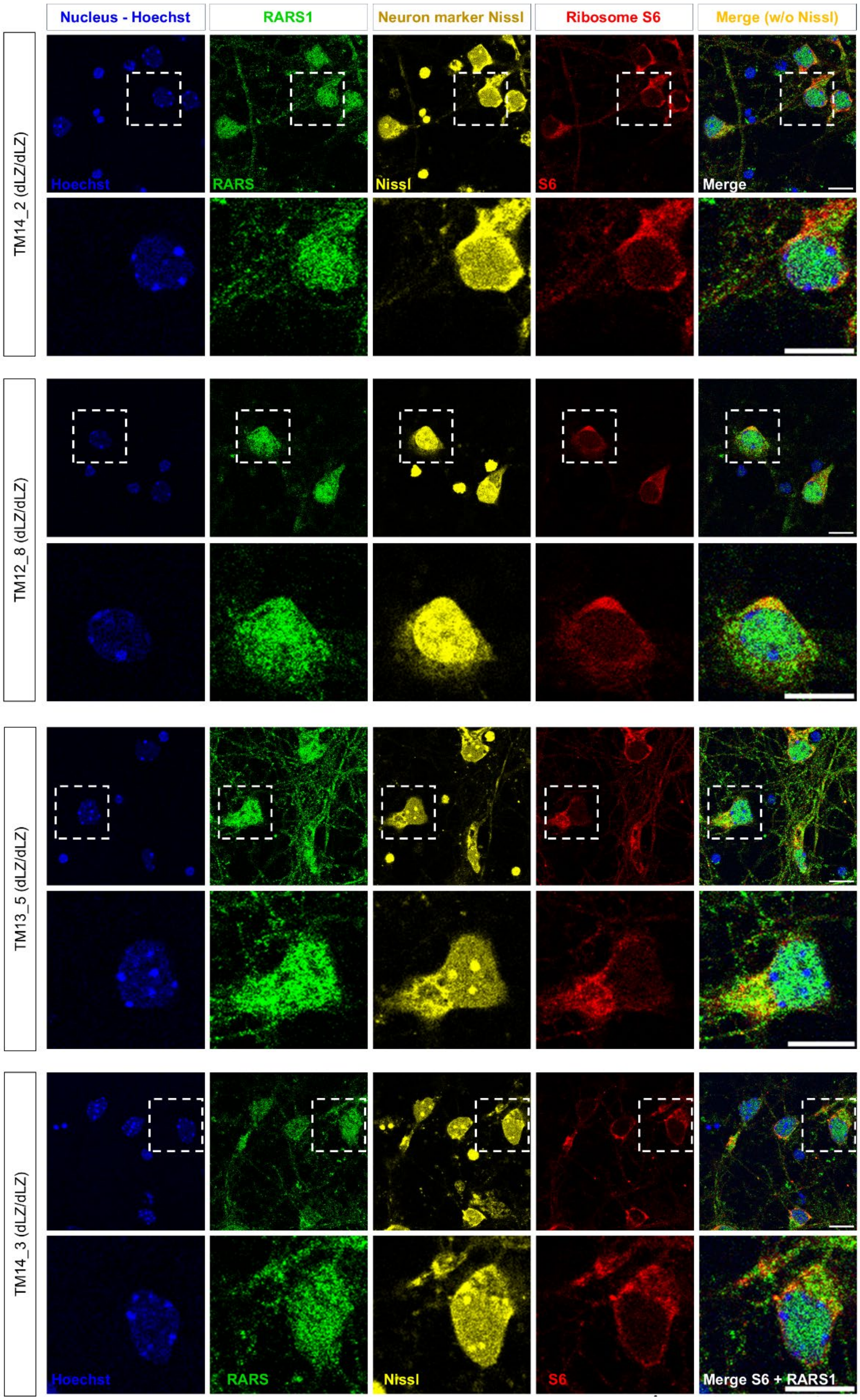

Nucleus - Hoechst
RARS1
Neuron marker Nissl
Ribosome S6
Merge (w/o Nissl)
TM14_2 (dLZ/dLZ)
TM12_8 (dLZ/dLZ)
TM13_5 (dLZ/dLZ)
TM14_3 (dLZ/dLZ)
Hoechst
RARS
Nissl
S6
Merge
Merge S6 + RARS1

**Extended Data Fig. 6: Exemplary images of RARS1 and S6 in primary neural cells and reduction of RARS1 and S6 in non-neuronal cells compared to neurons.**

A) Quantification of fluorescence intensity in neuronal cells compared to other cells. Both RARS1 and S6 were significantly higher expressed in neuronal cells. n=20, 16. n=20, 16. Unpaired, two-tailed t test, ****p<0.0001. B) Manders coefficient indicating colocalization of S6 relative to RARS1 in neuronal cell bodies. n=25, 28. Unpaired, two-tailed t test, ****p<0.0001. C) Representative images of isolated, cultured, and differentiated E14.5 cortical precursors. RARS1 and S6 were visualized using immunofluorescence. Hoechst was used to identify nuclei, indicating cell bodies. Bar=10 μm in the overview image, 5 μm in the zoomed in image. Cells isolated from 3 wt and 5 dLZ/dLZ embryos were assessed in total.

A-C) wt/wt: wt RARS1, systemic inclusion of RARS1 exon 2. dLZ/dLZ: RARS1 delta leucine zipper, systemic exclusion of RARS1 exon 2. Mice originated from *Sox2*-Cre x Rars1$^{loxP/loxP}$ and were bred as wt/dLZ animals following F1. Replicates are individual embryos. Sex was determined but results were not stratified by sex.

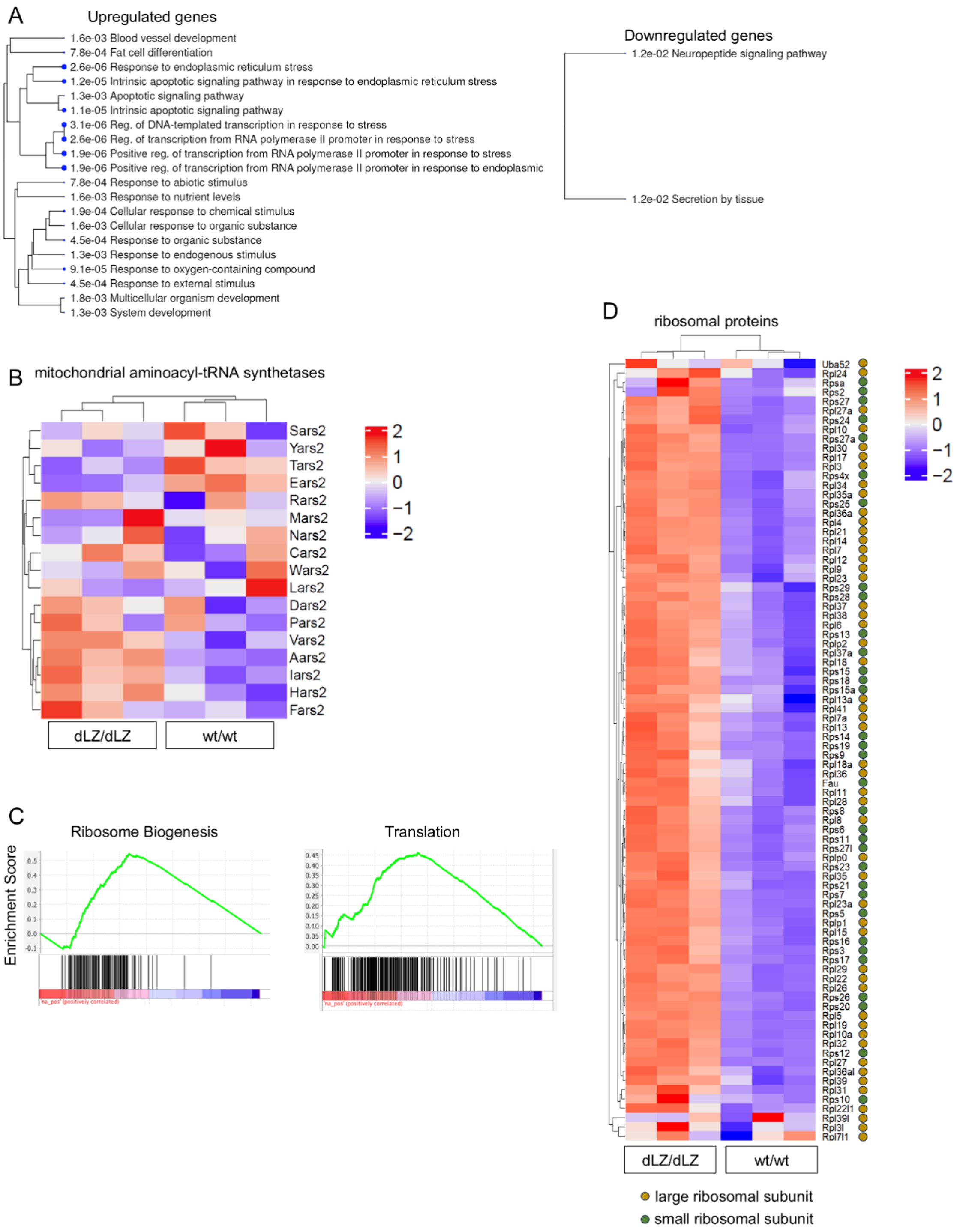
A
Upregulated genes
1.6e-03 Blood vessel development
7.8e-04 Fat cell differentiation
2.6e-06 Response to endoplasmic reticulum stress
1.2e-05 Intrinsic apoptotic signaling pathway in response to endoplasmic reticulum stress
1.3e-03 Apoptotic signaling pathway
1.1e-05 Intrinsic apoptotic signaling pathway
3.1e-06 Reg. of DNA-templated transcription in response to stress
2.6e-06 Reg. of transcription from RNA polymerase II promoter in response to stress
1.9e-06 Positive reg. of transcription from RNA polymerase II promoter in response to stress
1.9e-06 Positive reg. of transcription from RNA polymerase II promoter in response to endoplasmic
7.8e-04 Response to abiotic stimulus
1.6e-03 Response to nutrient levels
1.9e-04 Cellular response to chemical stimulus
1.6e-03 Cellular response to organic substance
4.5e-04 Response to organic substance
1.3e-03 Response to endogenous stimulus
9.1e-05 Response to oxygen-containing compound
4.5e-04 Response to external stimulus
1.8e-03 Multicellular organism development
1.3e-03 System development
Downregulated genes
1.2e-02 Neuropeptide signaling pathway
1.2e-02 Secretion by tissue
B
mitochondrial aminoacyl-tRNA synthetases
Sars2
Yars2
Tars2
Ears2
Rars2
Mars2
Nars2
Cars2
Wars2
Lars2
Dars2
Pars2
Vars2
Aars2
Iars2
Hars2
Fars2
2
1
0
−1
−2
dLZ/dLZ
wt/wt
C
Ribosome Biogenesis
Translation
Enrichment Score
D
ribosomal proteins
Uba52
Rpl24
Rpsa
Rps2
Rps27
Rpl27a
Rps24
Rpl10
Rps27a
Rpl30
Rpl17
Rpl3
Rps4x
Rpl34
Rpl35a
Rps25
Rpl36a
Rpl4
Rpl21
Rpl14
Rpl7
Rpl12
Rpl9
Rpl23
Rps29
Rps28
Rpl37
Rpl38
Rpl6
Rps13
Rplp2
Rpl37a
Rpl18
Rps15
Rps18
Rps15a
Rpl13a
Rpl41
Rpl7a
Rpl13
Rps14
Rps19
Rps9
Rpl18a
Rpl36
Fau
Rpl11
Rpl28
Rps8
Rpl8
Rps6
Rps11
Rps27l
Rplp0
Rps23
Rpl35
Rps21
Rps7
Rpl23a
Rps5
Rplp1
Rpl15
Rps16
Rps3
Rps17
Rpl29
Rpl22
Rpl26
Rps26
Rps20
Rpl5
Rpl19
Rpl10a
Rpl32
Rps12
Rpl27
Rpl36al
Rpl39
Rpl31
Rps10
Rpl22l1
Rpl39l
Rpl3l
Rpl7l1
2
1
0
−1
−2
dLZ/dLZ
wt/wt
large ribosomal subunit
small ribosomal subunit

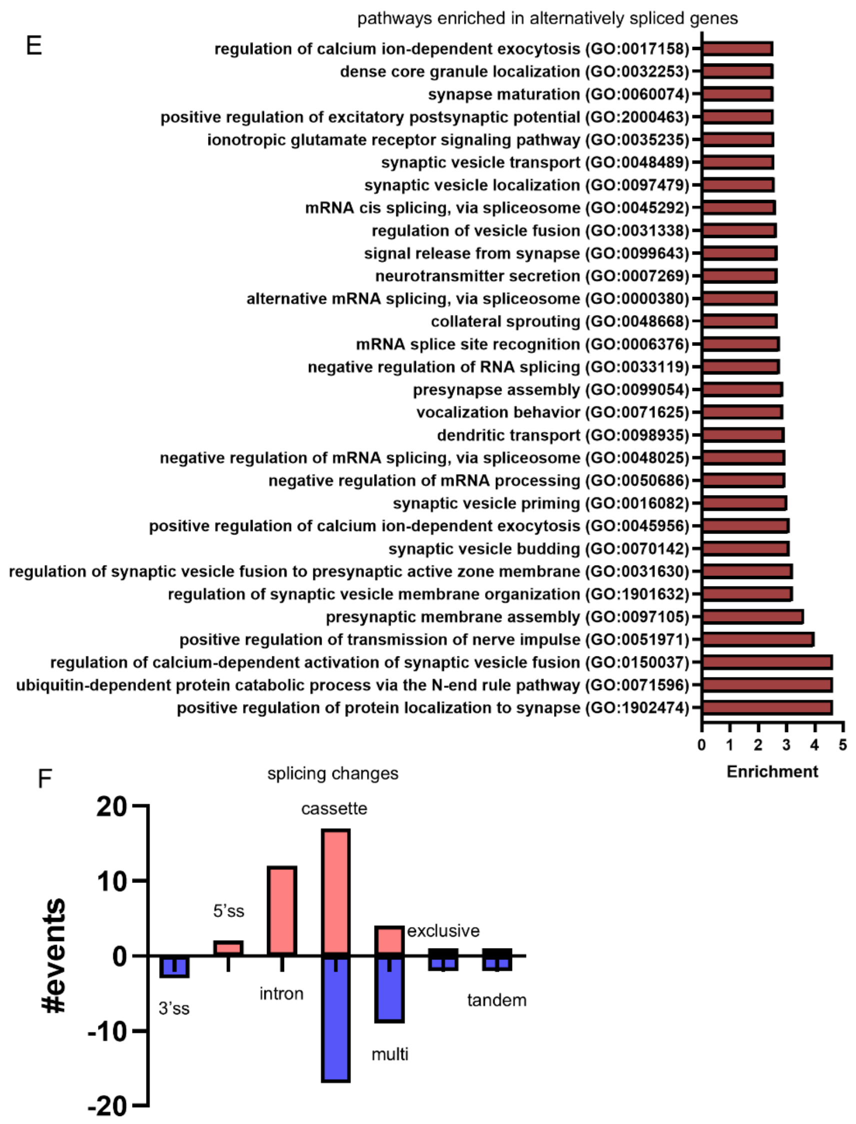
E
pathways enriched in alternatively spliced genes
regulation of calcium ion-dependent exocytosis (GO:0017158)
dense core granule localization (GO:0032253)
synapse maturation (GO:0060074)
positive regulation of excitatory postsynaptic potential (GO:2000463)
ionotropic glutamate receptor signaling pathway (GO:0035235)
synaptic vesicle transport (GO:0048489)
synaptic vesicle localization (GO:0097479)
mRNA cis splicing, via spliceosome (GO:0045292)
regulation of vesicle fusion (GO:0031338)
signal release from synapse (GO:0099643)
neurotransmitter secretion (GO:0007269)
alternative mRNA splicing, via spliceosome (GO:0000380)
collateral sprouting (GO:0048668)
mRNA splice site recognition (GO:0006376)
negative regulation of RNA splicing (GO:0033119)
presynapse assembly (GO:0099054)
vocalization behavior (GO:0071625)
dendritic transport (GO:0098935)
negative regulation of mRNA splicing, via spliceosome (GO:0048025)
negative regulation of mRNA processing (GO:0050686)
synaptic vesicle priming (GO:0016082)
positive regulation of calcium ion-dependent exocytosis (GO:0045956)
synaptic vesicle budding (GO:0070142)
regulation of synaptic vesicle fusion to presynaptic active zone membrane (GO:0031630)
regulation of synaptic vesicle membrane organization (GO:1901632)
presynaptic membrane assembly (GO:0097105)
positive regulation of transmission of nerve impulse (GO:0051971)
regulation of calcium-dependent activation of synaptic vesicle fusion (GO:0150037)
ubiquitin-dependent protein catabolic process via the N-end rule pathway (GO:0071596)
positive regulation of protein localization to synapse (GO:1902474)
0 1 2 3 4 5
Enrichment
F
splicing changes
20
10
0
-10
-20
#events
cassette
5'ss
exclusive
intron
3'ss
multi
tandem

**Extended Data Fig. 7: Remodelling of the transcriptome upon RARS1 exclusion from the multisynthetase complex.**

A) Enrichment plot based on ranked gene lists of differentially expressed genes (padj < 0.05) suggested the downregulation of genes associated with axon ensheathment, which can be indicative of myelination defects, and genes associated with mRNA translation. Genes associated with DNA transcription and external stimulus were enriched among upregulated genes in dLZ/dLZ forebrains compared to wt/wt. Genes associated with neuropeptide synthesis and secretion were enriched among downregulated genes in dLZ/dLZ forebrains compared to wt/wt. B) Heatmap of mitochondrial aminoacyl-tRNA synthetases showed no clear pattern. C) Enrichment of genes associated with ribosome biogenesis. D) Heatmap of genes encoding ribosomal proteins of both the small and large subunit showed consistent upregulation in dLZ/dLZ forebrains compared to wt/wt. D) Genes associated with mRNA processing and synaptic function were enriched among differentially spliced genes in dLZ/dLZ forebrains compared to wt. F) Overview of changes stratified by splicing event types.

A-E) wt/wt: wildtype RARS1, *Emx1*-Cre+ x Rars1$^{wt/wt}$. dLZ/dLZ: RARS1 delta leucine zipper, *Emx1*-Cre+ x Rars1$^{loxP/loxP}$. Replicates are individual animals. All experiments were performed in 2 months old male mice and are based on an RNAseq data set obtained with n=3, 3.

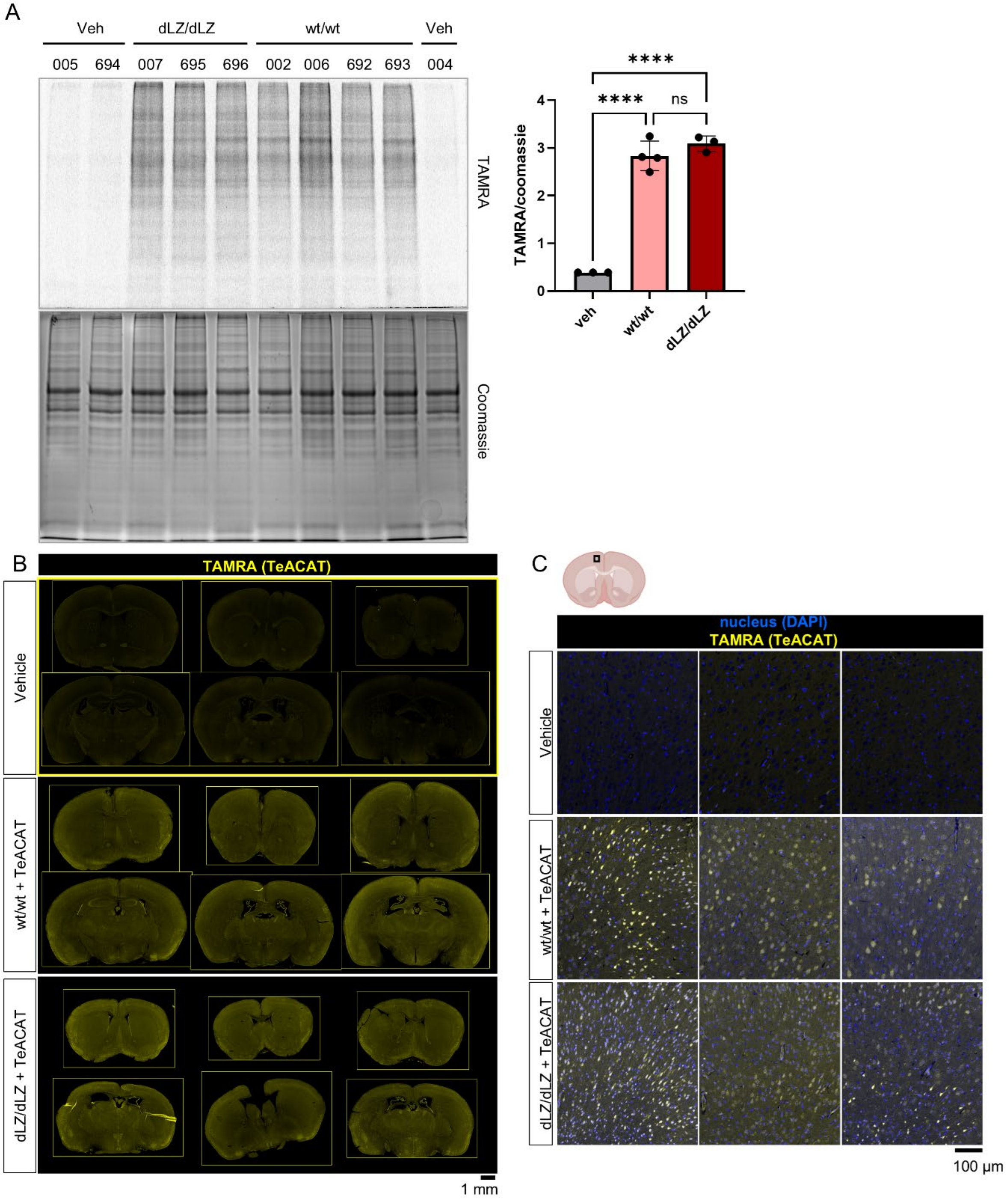
A
Veh
dLZ/dLZ
wt/wt
Veh
005
694
007
695
696
002
006
692
693
004
TAMRA
Coomassie
****
****
ns
TAMRA/coomassie
4
3
2
1
0
veh
wt/wt
dLZ/dLZ
B
TAMRA (TeACAT)
Vehicle
wt/wt + TeACAT
dLZ/dLZ + TeACAT
1 mm
C
nucleus (DAPI)
TAMRA (TeACAT)
Vehicle
wt/wt + TeACAT
dLZ/dLZ + TeACAT
100 µm

D

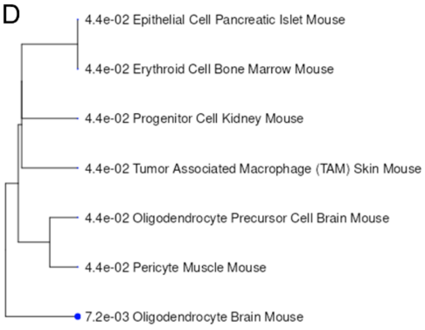


H

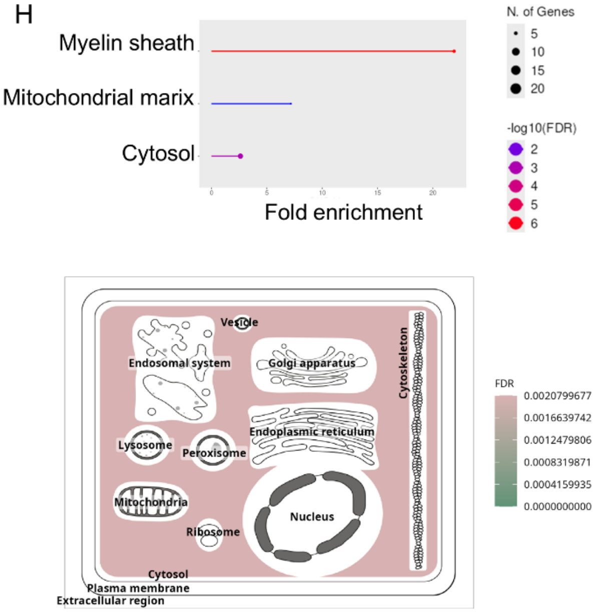


E

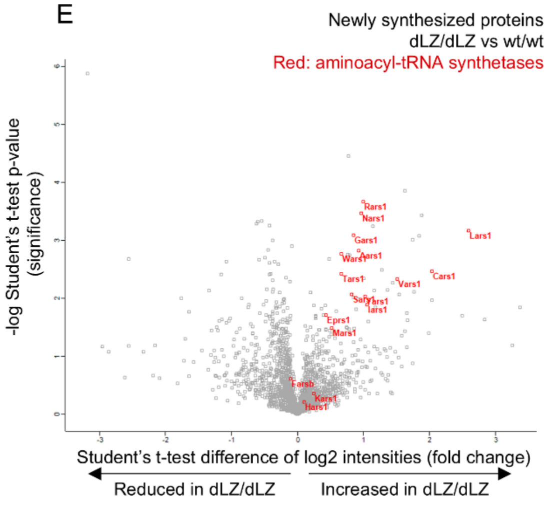


F

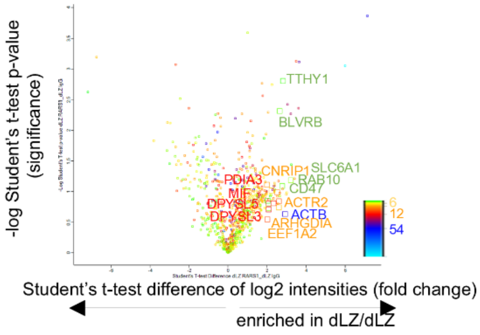


G Subcellular localization of proteins enriched in wt/wt:

| | | | | | | | | |
|---|---|---|---|---|---|---|---|---|
| Cnp | Cnrip1 | Slc25a4 | Rab3a | Dusp3 | Rtn1 | Plp1 | Septin8 | Crym |
| Luc7l2 | **Lars1** | **Aimp3** | **Kars1** | Nefm | **Dars1** | Nefl | **Qars1** | **Eprs1** |
| Aimp1 | Diras2 | **Iars1** | **Tars3** | Aimp2 | **Mars1** | | | |

Subcellular localization of proteins enriched in dLZ/dLZ:

| | | | | | | | | |
|---|---|---|---|---|---|---|---|---|
| **Yars1** | H3-3a | Ddx1 | **Aars1** | Nsfl1c | **Gars1** | Lonp1 | Hdhd2 | **Nars1** |
| **Wars1** | Poldip3 | Psph | Slc7a5 | Phgdh | Pdcd5 | Cyb5r1 | Srsf1 | Hdgf |
| Slc3a2 | Hbb-b1 | Ndrg2 | Epb41l2 | Basp1 | St13 | Etfa | Hsp90ab1 | Cox4i1 |
| Ncam1 | Arpc1a | Wdr13 | Map2 | Atp1b1 | | | | |

Cytoplasmic
Plasma membrane/myelin sheath
Nuclear
Nuclear & cytoplasmic
Mitochondrial

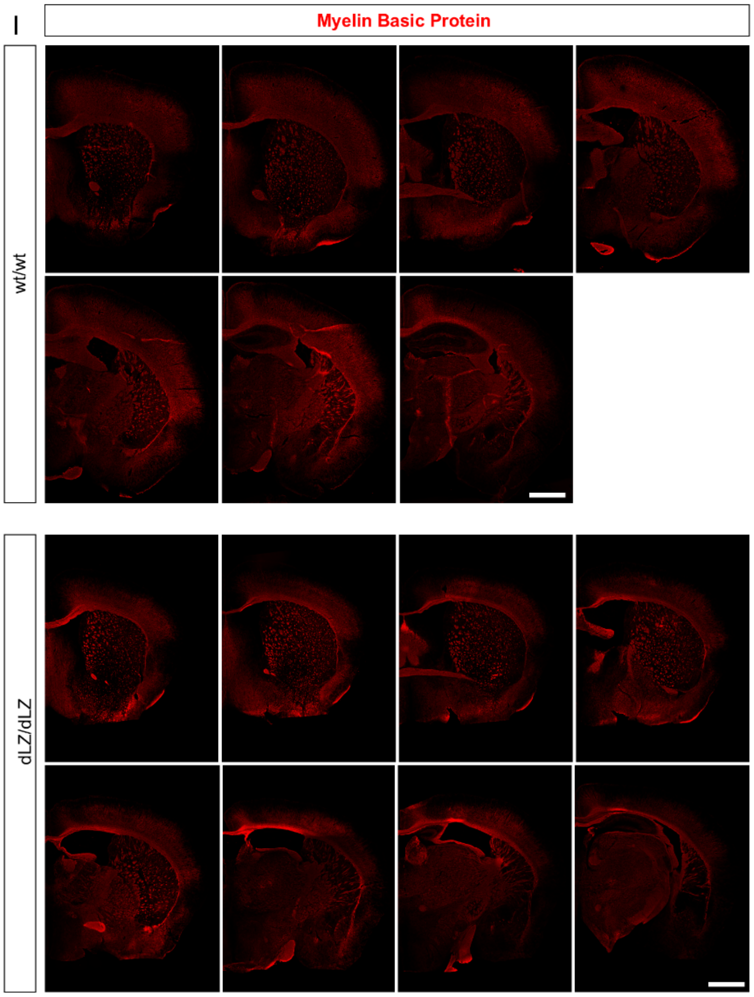
I
Myelin Basic Protein
wt/wt
dLZ/dLZ

**Extended Data Fig. 8: Tellurophene-Alkyne Cycloaddition-mediated Amino acid Tagging (TeACAT) in mouse brains and analysis of the RARS1 interactome.**

A) SDS-PAGE of olfactory bulbs lysates following TePhe injection and visualization of newly synthesized proteins using TeACAT. TAMRA fluorescence in the gels to quantify TePhe incorporation and Coomassie to quantify total protein in the lysates. OSTAC intensity normalized to Coomassie total protein levels. One-way ANOVA, multiple comparisons, ****$p < 0.0001$, ****$p < 0.0001$, ns $p > 0.05$. B) TeACAT-based detection of TePhe incorporation marked by TAMRA into newly synthesized proteins in FFPE sections. Bulk protein synthesis seems comparable between wt/wt and dLZ/dLZ forebrains. Scale bar = 1 mm. C) Higher magnification images of TeACAT sections shows protein synthesis in cell bodies and cellular projections. Bar = 100 µm. D) Cell type analysis of newly synthesized proteins with reduced protein synthesis highlights that proteins found in oligodendrocytes are especially affected by RARS1 exclusion from the multisynthetase complex. E) Aminoacyl-tRNA synthetase protein synthesis was increased in dLZ/dLZ mice. Volcano plot of proteins identified by mass spectrometry following OSTAC labelling, biotin tagging and streptavidin pulldown. *Y*-axis: significance (*p* value). *X*-axis: difference (fold change). Comparison between proteins enriched in wt and dLZ/dLZ lysates. Proteins enriched more significantly in wt pulldowns than dLZ RARS1 appear on the *left side* of the volcano plot and proteins enriched more significantly in dLZ/dLZ lysates than wt appear on the *right* side of the plot. Student's t-test. F) Volcano plot of proteins identified by mass spectrometry following coimmunoprecipitation of RARS1 in dLZ/dLZ mice, highlighting neural proteins. New *Y*-axis: significance (*p* value). *X*-axis: difference (fold change). Color coding denotes the number of peptides per protein. RARS1 dLZ/dLZ interactome (anti-RARS1 antibody pulldown compared with IgG control pulldown in dLZ/dLZ brain lysates). Color coding denotes the number of peptides. n = 3, 3. Student's t-test. G) List of enriched proteins in wt/wt and dLZ/dLZ RARS1 interactomes, colored by cellular localization. H) Gene enrichment analysis of proteins only found within the RARS1 dLZ/dLZ interactome and visualization of protein localization within the cell. I) Immunofluorescence of MBP.

A-I) wt/wt: wildtype RARS1, *Emx1*-Cre+ x Rars1$^{wt/wt}$. dLZ/dLZ: RARS1 delta leucine zipper, *Emx1*-Cre+ x Rars1$^{loxP/loxP}$. Replicates are individual animals. All experiments were performed in 2 months old male mice

A-F) Based on a TeACAT data set obtained with n=3, 3.

F-H) Based on a RARS1 interactome data set obtained with n=3, 3.

**Supplementary Table legends:**

**Supplementary Table 1:** wt/wt RARS1 interactome in mouse forebrains.

**Supplementary Table 2:** dLZ/dLZ RARS1 interactome in mouse forebrains.

**Supplementary Table 3:** Comparison of wt/wt RARS1 interactomes with dLZ/dLZ RARS1 interactomes.

**Supplementary Table 4:** Differential gene expression in mouse forebrains by RNAseq.

**Supplementary Table 5:** Differential mRNA splicing in mouse forebrains.

**Supplementary Table 6:** Newly synthesized proteins enriched from wt/wt forebrains with TeACAT-MS compared to vehicle controls.

**Supplementary Table 7:** Newly synthesized proteins enriched in dLZ/dLZ forebrains through TeACAT-MS compared to vehicle controls.

**Supplementary Table 8:** Differentially newly synthesized proteins in wt/wt vs dLZ/dLZ forebrains.

**Supplementary Table 9:** Antibodies used in this study.
